# Characterization of a spectrally diverse set of fluorescent proteins as FRET acceptors for mTurquoise2

**DOI:** 10.1101/156448

**Authors:** Marieke Mastop, Daphne S. Bindels, Nathan C. Shaner, Marten Postma, Theodorus W. J. Gadella, Joachim Goedhart

## Abstract

Genetically encoded Förster Resonance Energy Transfer (FRET) based biosensors report on changes in biochemical states in single living cells. The performance of biosensors depends on their brightness and dynamic range, which are dependent on the characteristics of the fluorescent proteins that are employed. Cyan fluorescent protein (CFP) is frequently combined with yellow fluorescent protein (YFP) as FRET pair in biosensors. However, current YFPs are prone to photobleaching and pH changes. In addition, more efficient acceptors may yield biosensors that have higher contrast. In this study, we evaluated the properties of a diverse set of acceptor fluorescent proteins in combination with the optimized CFP variant mTurquoise2 as the donor. To determine the theoretical performance of acceptors, the Förster radius was determined. The practical performance was determined by measuring FRET efficiency and photostability of tandem fusion proteins in mammalian cells. Our results show that mNeonGreen is the most efficient acceptor for mTurquoise2 and that the photostability is better than SYFP2. The non-fluorescent YFP variant sREACh is an efficient acceptor, which is useful in lifetime-based FRET experiments. Among the orange and red fluorescent proteins, mChery and mScarlet-I are the best performing acceptors. Several new pairs were applied in a multimolecular FRET based sensor for detecting activation of a heterotrimeric G-protein by G-protein coupled receptors. The sensor with mScarlet-I as acceptor and mTurquoise2 as donor shows a higher dynamic range in ratiometric FRET imaging experiments and less variability than with mCherry as acceptor, due to the high quantum yield and efficient maturation of mScarlet-I. Overall, the sensor with mNeonGreen as acceptor and mTurquoise2 as donor showed the highest dynamic range in ratiometric FRET imaging experiments with the G-protein sensor.

## Introduction

Fluorescent proteins derived from jellyfish and corals are fluorescent probes that are entirely genetically encoded and do not require a co-factor ^1,2^. These probes are important tools for fluorescence imaging of cellular processes ^3,4^ A specific application of fluorescent proteins is their use in Förster resonance energy transfer (FRET) studies ^5–8^. FRET is the radiationless transfer of energy from an excited donor to a nearby acceptor. The FRET efficiency depends on several parameters, including the quantum yield of the donor, the extinction coefficient of the acceptor and the spectral overlap of donor emission and acceptor absorbance ^9,10^. The aforementioned parameters determine the Förster distance, *R*_*0*_, which is the distance between donor and acceptor that will result in 50% FRET ^11,12^.

FRET can be used to determine the interaction between biomolecules and is also the basis for so-called biosensors. Biosensors are designed to report on chemical states and can be used to measure concentrations of ions or small molecules, phosphorylation of peptides or the nucleotide loading state of a protein ^13,14^. The performance of FRET based biosensors depends on their brightness and dynamic range, which are highly dependent on the characteristics of the applied fluorescent proteins ^15–17^. Both FRET efficiency and brightness depend on extinction coefficient and quantum yield and therefore a general recommendation is to use the brightest fluorescent proteins available ^17^ For FRET imaging in living cells, several other parameters should be considered including maturation, photostability, oligomeric state and sensitivity to environmental changes ^8,12,17^.The maturation is a critical factor for effective brightness of a fluorescent protein and for efficient FRET ^6^. The maturation efficiency is the fraction of produced protein that results in a correctly folded protein with a functional, fluorescent chromophore. Ideally, the maturation of a fluorescent protein approaches 100%. When a protein incorrectly folds or does not form a correct chromophore, the FRET pair will lack a functional donor or acceptor and this will prevent FRET, thereby diluting the number of functional FRET pairs and decreasing the dynamic range ^17^. In *Aequorea victoria* derived fluorescent proteins, amino acid residues ^65–67^ of the folded protein undergo several chemical reactions necessary for chromophore formation, including cyclization, oxidation and dehydration ^2^ Characteristics of residues in the vicinity of the chromophore can influence the efficiency of protein folding and chromophore formation. Mutations leading to more efficient chromophore formation (F64L, V68L) or protein folding (S72A, V163A, S175G) were identified ^18–23^However, other mutations may lead to inefficient or slow maturation, resulting in dim fluorescence and only a small fraction of fluorescent cells ^24,25^ In red fluorescent proteins (RFPs), the maturation process is more complex. After the cyclization and oxidation steps the chromophore can be dehydrogenated in two alternative ways. One leads to a blue fluorescent intermediate that upon another oxidation step results in a mature RFP, while the other leads to a non-reversible GFP form ^26^. Thus in the case of RFPs, inefficient or slow maturation may result in substantial green or blue fluorescence next to dim and inefficient RFP expression, hindering their use in multi-color labeling experiments ^1^.

Furthermore, it is important that the fluorescent proteins used in biosensors are not sensitive to environmental changes other than the one you want to measure. For example, yellow fluorescent protein (YFP) variants are sensitive to halide concentrations, but this problem was addressed by mutagenesis resulting in YFP variants Citrine and Venus ^24,27,28^. Next to sensitivity for halides, pH sensitivity could also affect the absorbance and hence FRET efficiency (mainly in GFP, YFP and mOrange). The pH sensitivity is dependent on the pKa of a fluorescent protein and depending on the acidity of the experimental environment, this characteristic should be taken into account when choosing or constructing a biosensor.

In nature, fluorescent proteins usually exist as dimers or tetramers ^2,29,30^. It is important that fluorescent proteins that are tagged to proteins of interest are not oligomerizing, because this can lead to impaired functioning and/or localization of the protein of interest and it can lead to false positives in interaction studies^8,31^. The latter issue is more critical for intermolecular sensors as compared to intramolecular sensors. In fact, a weak tendency of heterodimerization can be beneficial for FRET contrast for unimolecular sensors ^32–34^. Monomeric variants of *Aequorea victorea* fluorescent proteins were obtained by replacing hydrophobic residues at the dimer interface with positively charged residues (A206K, L221K, or F223R) ^31^. The engineering of bright, monomeric RFP variants is more difficult, since mutations disrupting dimer interfaces also affect other characteristics such as the quantum yield ^35^. A recent engineering effort has resulted in a truly monomeric red fluorescent protein, mScarlet-I, with good maturation. Because of its relatively high quantum yield, the level of sensitized emission surpasses that of mCherry in a FRET pair ^36^.The monomeric nature of fluorescent proteins is often analyzed via *in vitro* ultra centrifugation or gel filtration of purified proteins ^30,31,37^ and this is not a good predictor for the tendency to dimerize in living cells. Costantini et al. developed an *in vivo* dimerization assay in which fluorescent proteins are fused to an endoplasmic reticulum (ER) signal anchor membrane protein (CytERM). Homo-oligomerization of this CytERM-FP with the same construct in opposing membranes causes the formation of organized smooth ER (OSER) structures, which can be quantitatively evaluated in this OSER assay ^38,39^. Recently, Cranfill et al. assessed the oligomeric state of a large number of fluorescent proteins in cells using the OSER assay ^40^, providing a useful guide in choosing fluorescent proteins for certain applications.

FRET based sensors are mostly used in dynamic systems that are examined by timelapse imaging and therefore, photostability is an important characteristic. During timelapse imaging, it is crucial that only the actual changes in FRET are reported, since differences in photobleaching characteristics between the fluorescent proteins in a sensor will result in false FRET changes, complicating data analysis. Since FRET by itself changes photobleaching rates ^41^, changes in FRET will result in altered photobleaching kinetics. Hence, the photobleaching rate may change during a timelapse experiment and therefore it is close to impossible to correct for photobleaching. Consequently, it is important to choose photostable fluorescent proteins, enabling FRET imaging with little photobleaching.

The photostability of fluorescent proteins is only poorly understood. The photostability differs even between fluorescent proteins with very similar optical properties ^42,43^. The β-barrel around the chromophore protects the chromophore against oxidative damage so perhaps slight changes in the β-barrel architecture account for these differences ^44–47^. Recently, it was reported that many fluorescent proteins show supralinear photobleaching. Consequently, if the excitation light power doubles, the photobleaching rate increases with a factor of more than two ^40^ Therefore, photostability depends on the illumination power this should be taken into consideration when choosing fluorescent proteins for a FRET pair. In addition, photochromic behavior and photoconversion can also drastically change the intensity of a fluorophore over time and therefore should be evaluated as well ^15,36^.

The photostability of fluorescent proteins is usually determined at the excitation wavelength that is close to the absorbance maximum ^40,43^. However, in FRET experiments, either FLIM or ratio-imaging, FRET acceptors are usually excited far from their absorbance maximum. In addition, they receive energy from the excited donor. Exactly, how these different modes and wavelengths of excitation affect the photostability of acceptor fluorophores, and consequently the FRET pair, has not been thoroughly investigated.

At the moment cyan fluorescent protein (CFP) or teal fluorescent protein (TFP) combined with yellow fluorescent protein (YFP) is the most frequently used as FRET pair in biosensors ^48–51^. The CFP variant mTurquoise2 is an attractive FRET donor because of its high quantum yield (of 93%), monomeric behavior and good photostability ^40,42^. As for acceptors, optimized variants of YFP: mCitrine, mVenus, YPet and SYFP2 (mVenus-L68V), are reported ^18,24,27, 34^. These YFPs exhibit a high extinction coefficient, optimized folding, a large spectral overlap with the emission spectrum of mTurquoise2 and a good quantum yield. However, current YFPs lack photostability and pH-stability. In addition, acceptors that provide an even higher FRET efficiency might yield biosensors that have improved contrast. Therefore, in this study, we evaluated the properties of a diverse set of acceptor proteins in combination with mTurquoise2 as donor.

Many studies have reported improvements of FRET sensors by changing the distance between the fluorescent proteins, varying linker length ^52–54^ and/or composition ^55^ or changing the relative orientation of the fluorescent proteins by using circular permuted fluorescent protein variants ^48,51^. Recently, it was reported that even the order of fluorescent proteins in a sensor alters its dynamic range ^56^.

Here, we aspired to examine which of the current bright fluorescent proteins would have favorable properties for FRET-based imaging, not taking into account linkers and relative orientation. To this end, we evaluated the FRET efficiencies of FRET pairs consisting of mTurquoise2 as donor and acceptors varying from green to far-red. The Förster distance was determined for every pair, followed by experimental determination of FRET efficiencies of tandem fluorescent protein constructs in living cells. The FRET efficiencies were determined by fluorescence lifetime imaging (FLIM) and spectral imaging microscopy (SPIM) of tandem fusions. In addition, the photostability under FRET conditions was determined. The most promising pairs were applied in a FRET based biosensor for heterotrimeric G-protein activation.

## Results

### Absorption and emission spectra of purified fluorescent proteins

Due to the long emission tail of mTurquoise2, fluorescent proteins red-shifted relative to mTurquoise2 are potentially efficient FRET acceptors. We selected a number of promising acceptor candidates based on two criteria: (i) reported monomeric, (ii) bright in their spectral class. The list of selected proteins covers the visible spectrum, with fluorescent protein emission colors ranging from green to far-red. To judge the theoretical quality of the FRET pairs, we determined the Förster radius (*R*_*0*_).

In order to do so, we purified a selection of fluorescent proteins and determined the absorbance and emission spectra. The absorbance and emission spectra of the proteins employed in this study are depicted in figure 1 and the spectral data is published elsewhere (http://doi.org/10.5281/zenodo.580169). We note that the absorbance spectra of all the fluorescent proteins, even the most red-shifted variant, mKate2, overlap with the emission of mTurquoise2.

**Figure 1.**
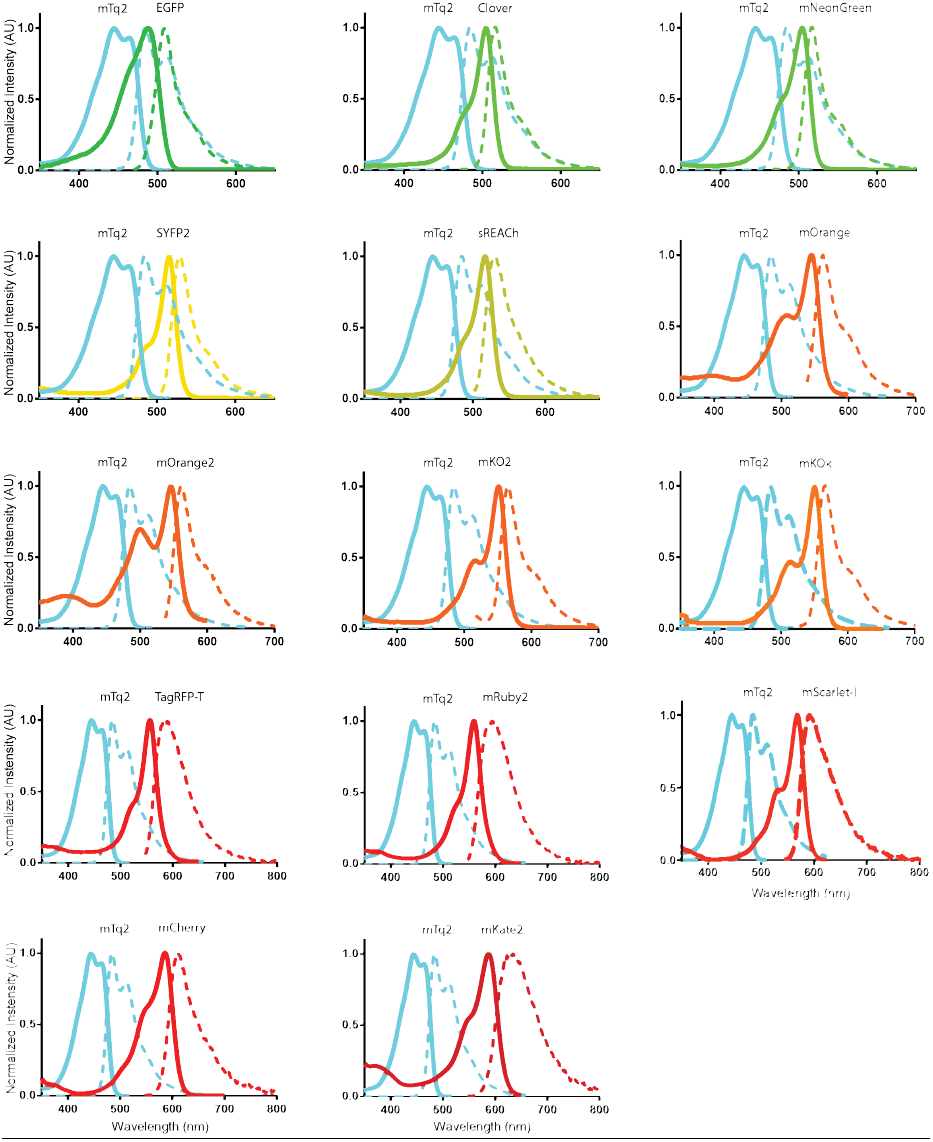
Absorption and emission spectra of the FRET pairs investigated in this study. The spectra were recorded from purified proteins and were normalized to their peak values. Solid lines indicate absorption spectra and dashed lines indicate emission spectra. All lines are colored according to the emission wavelength of the fluorescent protein. All spectra show the donor mTurquoise2 (mTq2) and the indicated acceptor. Data available at http://doi.org/10.5281/zenodo.580169.

Next, we determined the overlap integral, *J(λ)*, for mTq2 emission with the absorbance, based on the spectra that we acquired and the published extinction coefficient of the acceptors (table 1). The overlap integral was used to calculate the Förster radius *R*_*0*_, assuming a refractive index (*n*) of 1.33 and *κ*^*2*^of 2/3 ^11,15^. Of note, *n* and *κ*^*2*^are usually unknown in cells, but alternative *R*_*0*_ values can be calculated if *n* and *κ*^*2*^are known from the reported overlap integral.

**Table 1.** Spectroscopic parameters of fluorescent proteins employed in this study as acceptor for mTurquoise2. The overlap integral was determined from spectra acquired in this study and extinction coefficients taken from literature. The Föster radius was calculated from *J(λ)* and quantum yield of the donor (*QY*_*D*_) =0.93, *n*=1.33 and K^*2*^ = 2/3.

| FRET Acceptor | Absorption peak (nm) | Molar extinction coefficient ($M^{-1}cm^{-1}$) | $J(\lambda)*10^{15} M^{-1} cm^{-1} nm^4$ | $QY_A$ | $R_0$ (Å) |
| --- | --- | --- | --- | --- | --- |
| **EGFP** | 488 | 55000<sup>1</sup> | 1.53 | 0.6<sup>1</sup> | 55 |
| **Clover** | 505 | 111000<sup>2</sup> | 2.7 | 0.76<sup>2</sup> | 60 |
| **mNeonGreen** | 505 | 116000<sup>3</sup> | 3.15 | 0.8<sup>3</sup> | 62 |
| **SYFP2** | 515 | 101000<sup>4</sup> | 2.31 | 0.68<sup>4</sup> | 59 |
| **sREAcH** | 517 | 100000<sup>5</sup> | 2.53 | - | 59 |
| **mOrange** | 548 | 71000<sup>6</sup> | 2.19 | 0.69<sup>6</sup> | 58 |
| **mOrange2** | 549 | 58000<sup>7</sup> | 2.05 | 0.60<sup>7</sup> | 57 |
| **mKO2** | 551 | 63800<sup>8</sup> | 1.44 | 0.57<sup>8</sup> | 54 |
| **mKOκ** | 551 | 105000<sup>9</sup> | 2.08 | 0.61<sup>9</sup> | 58 |
| **TagRFP-T** | 557 | 81000<sup>7</sup> | 1.62 | 0.41<sup>7</sup> | 55 |
| **mRuby2** | 560 | 113000<sup>2</sup> | 2.12 | 0.38<sup>2</sup> | 58 |
| **mScarlet-I** | 569 | 102000<sup>10</sup> | 1.84 | 0.54<sup>10</sup> | 56 |
| **mCherry** | 587 | 72000<sup>6</sup> | 1.24 | 0.22<sup>6</sup> | 53 |
| **mKate2** | 588 | 62500<sup>11</sup> | 1.15 | 0.40<sup>11</sup> | 52 |

The values for the molar extinction coefficients, $\epsilon$ , were taken from literature: <sup>1</sup> ref. 22
<sup>2</sup> ref. 85, <sup>3</sup> ref. 63, <sup>4</sup> ref. 18, <sup>5</sup> ref. 69,86, <sup>6</sup> ref. 58, <sup>7</sup> ref. 43, <sup>8</sup> ref. 87, <sup>9</sup> ref. 68, <sup>10</sup> ref. 36, <sup>11</sup> ref. 70.

The calculated *R*_*0*_ values show a declining trend when the absorbance peak shifts to the red part of the spectrum. The standard cyan-yellow pair has a *R*_*0*_ of 59 Å. In theory, the best green acceptor is mNeonGreen with a *R*_*0*_ of 62 Å. The orange and red fluorescing fluorescent proteins with the highest *R*_*0*_ values are mKOk, mOrange and mRuby2 with a value of 58 Å.

In summary, from the Förster radii it can be concluded that the selected fluorescent proteins are promising as FRET acceptor since the *R*_*0*_ values are all above 50 Å.

### Fluorescence lifetime analysis of FRET pairs

The *R*_*0*_ values can be used as a theoretical measure for the quality of a FRET pair. However, it is important to evaluate the FRET pairs experimentally *in cyto*, to reveal cellular parameters that affect the FRET efficiency. To judge the quality of the FRET acceptors in cells, we constructed plasmids encoding fusion proteins incorporating mTurquoise2 as the donor and one of the candidate fluorescent proteins as the acceptor (figure 2). These plasmids were transfected in mammalian cells and fluorescence lifetime imaging microscopy (FLIM) was performed. In order to calculate the FRET efficiency, the donor lifetimes of cells in FRET and non-FRET conditions were measured. We used cells expressing untagged mTurquoise2 as non-FRET condition. These show a donor phase lifetime of 3.8ns, as reported before ^42^. The fusion constructs are used for FLIM measurements in FRET condition. All FRET pairs show a decrease in donor lifetime compared to untagged mTurquoise2 indicating that FRET occurred. For a complete overview of phase and modulation lifetime values and the FRET efficiency based on lifetime see table 2. We focused on the FRET efficiencies based on phase lifetime rather than modulation lifetime, because it shows a higher dynamic range meaning that differences in FRET efficiency will be more noticeable ^15^. The phase lifetimes are graphically depicted in figure 3. From figure 3 can be inferred that mNeonGreen shows the largest reduction in fluorescence lifetime and consequently the highest FRET efficiency *in cyto*. The other yellow-green acceptor fluorescent proteins, including the non-emitting variant sREACh, display lifetimes similar to the standard mTurquoise2-SYFP2 pair. Of note, standard SYFP2 shows high cell-to-cell variation compared to the other green and yellow acceptors. Among the orange acceptors, mKO*κ* shows the largest lifetime change, whereas mOrange and mOrange2 show only moderate changes in fluorescence lifetime and also display quite some cell-to-cell variability. The tandems that comprise red acceptors display similar lifetime reductions, with mRuby2 as the most efficient FRET acceptor. In summary, mNeonGreen shows the highest FRET efficiency and mKO*κ* stands out amongst the orange acceptors.

**Table 2.**
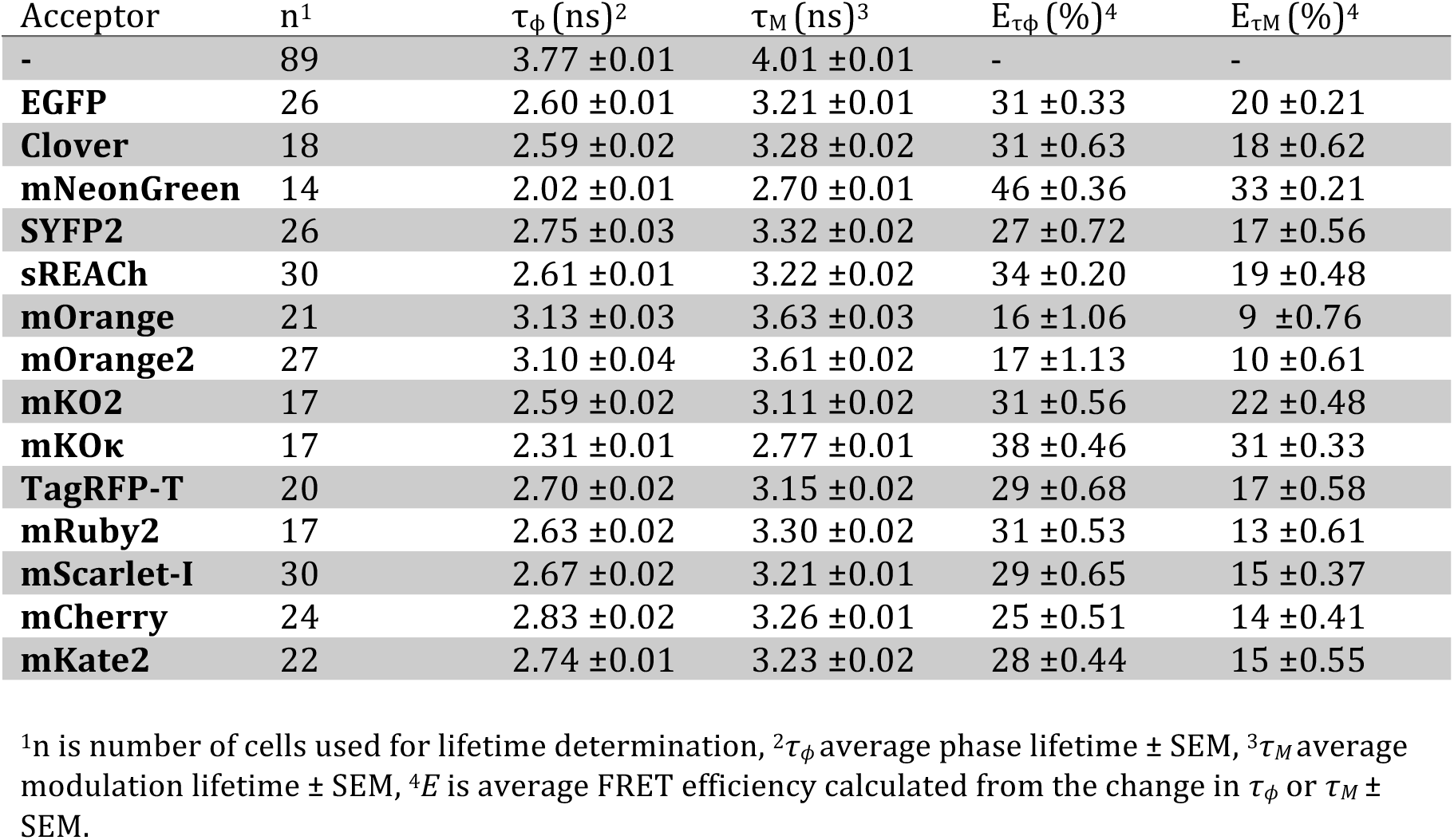
Fluorescence lifetime data of mTurquoise2 as FRET donor in a tandem construct with the different FRET acceptors and corresponding FRET efficiency (figure 3).

**Figure 2.**
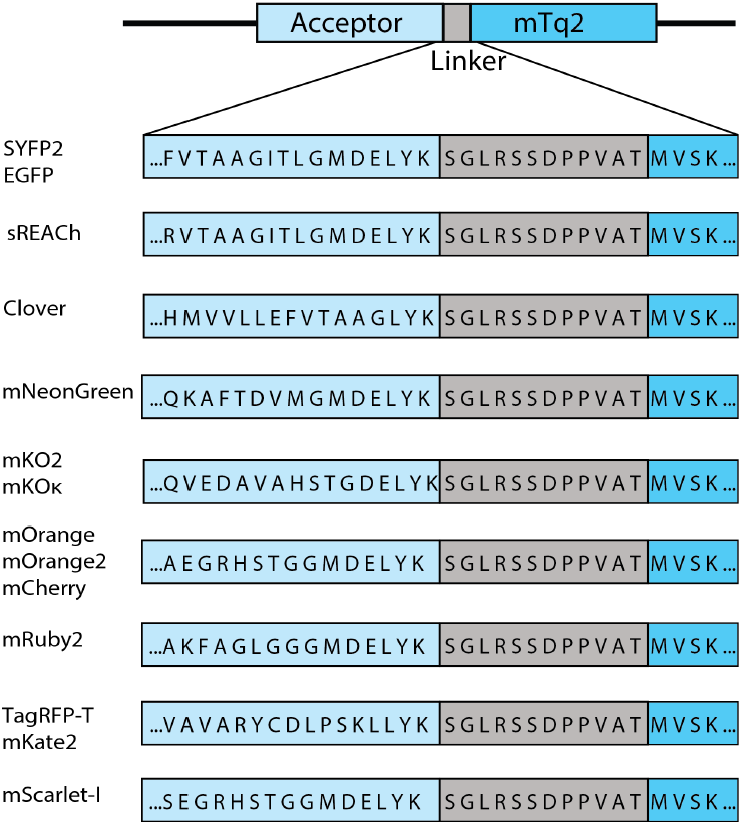
Schematic overview of the fusion constructs used in this study. The differences in the amino acid sequence of the C-termini of the acceptor fluorescent proteins are depicted. The size of the acceptors is 158 amino acids for mKO2 and mKO*κ*, 163 amino acids for mOrange, mOrange2, mScarlet-I and mCherry, 164 amino acids for Clover, 166 amino acids for mNeonGreen, TagRFP-T and mKate2, 168 amino acids for mRuby2 and 171 amino acids for EGFP, SYFP2 and sREACh. The acceptors are followed by a small linker, which is the same for each construct, separating it from the donor mTurquoise2 (mTq2).

**Figure 3.**
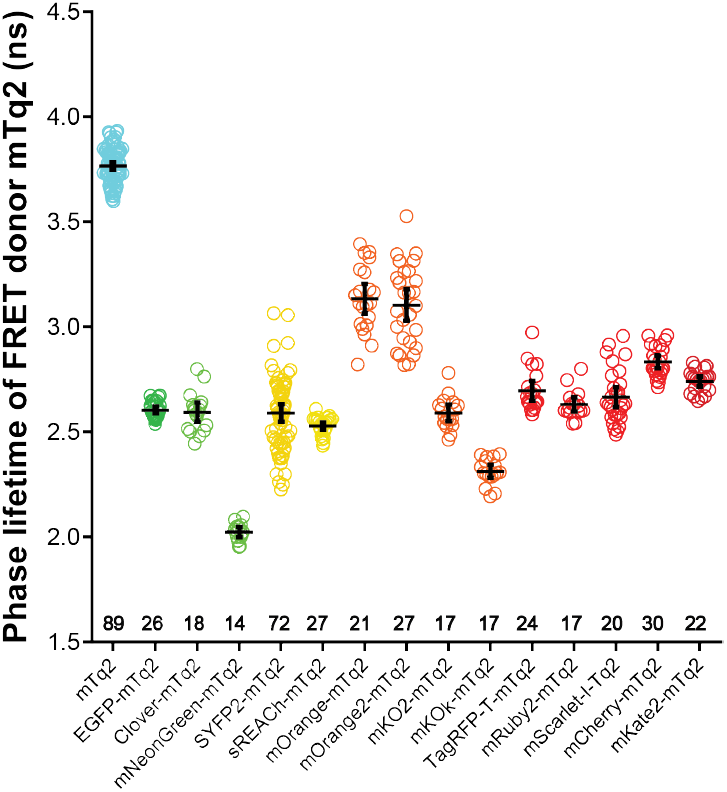
Fluorescence lifetime of the FRET donor mTurquoise2 fused to different FRET acceptors. The phase lifetime of mTurquoise2 (mTq2) when paired with different acceptors is depicted. As a reference the lifetime of untagged mTurquoise2 is shown. The dots indicate individual cells and the error bars show 95% confidence intervals. The number of cells imaged is mTq2 *n*=89, EGFP *n*=26, Clover *n*=18, mNeonGreen *n*=14, SYFP2 *n*=72, sREACh *n*=27, mOrange *n*=21, mOrange2 *n*=27, mKO2 *n*=17, mKO*κ n*=17, TagRFP-T *n*=20, mRuby2 *n*=17, mScarlet-I *n*=30, mCherry *n*=24, mKate2 *n*=22.

### Spectral imaging of FRET pairs

The FLIM data of FRET pairs gives insight in the importance of spectral overlap and extinction coefficient of the acceptor, while the quantum yield of the acceptor does not matter in FLIM measurements. Most of the currently applied biosensors are, however, analyzed by ratiometric imaging which relies, besides donor quenching, on sensitized emission ^8,57^. The sensitized emission depends on the FRET efficiency (spectral overlap and extinction coefficient) and the quantum yield of the acceptor. A higher sensitized emission results in a better contrast in ratiometric FRET imaging. To examine the amount of sensitized emission for each FRET pair, we acquired spectral images of single cells producing fusion proteins (figure 4). Corrected spectra were obtained by correcting for spectral sensitivity (tail of long-pass (LP) filter and camera). From these data, we isolated the pure sensitized emission component by unmixing the donor spectrum and the amount of direct acceptor excitation (figure 4).

**Figure 4.**
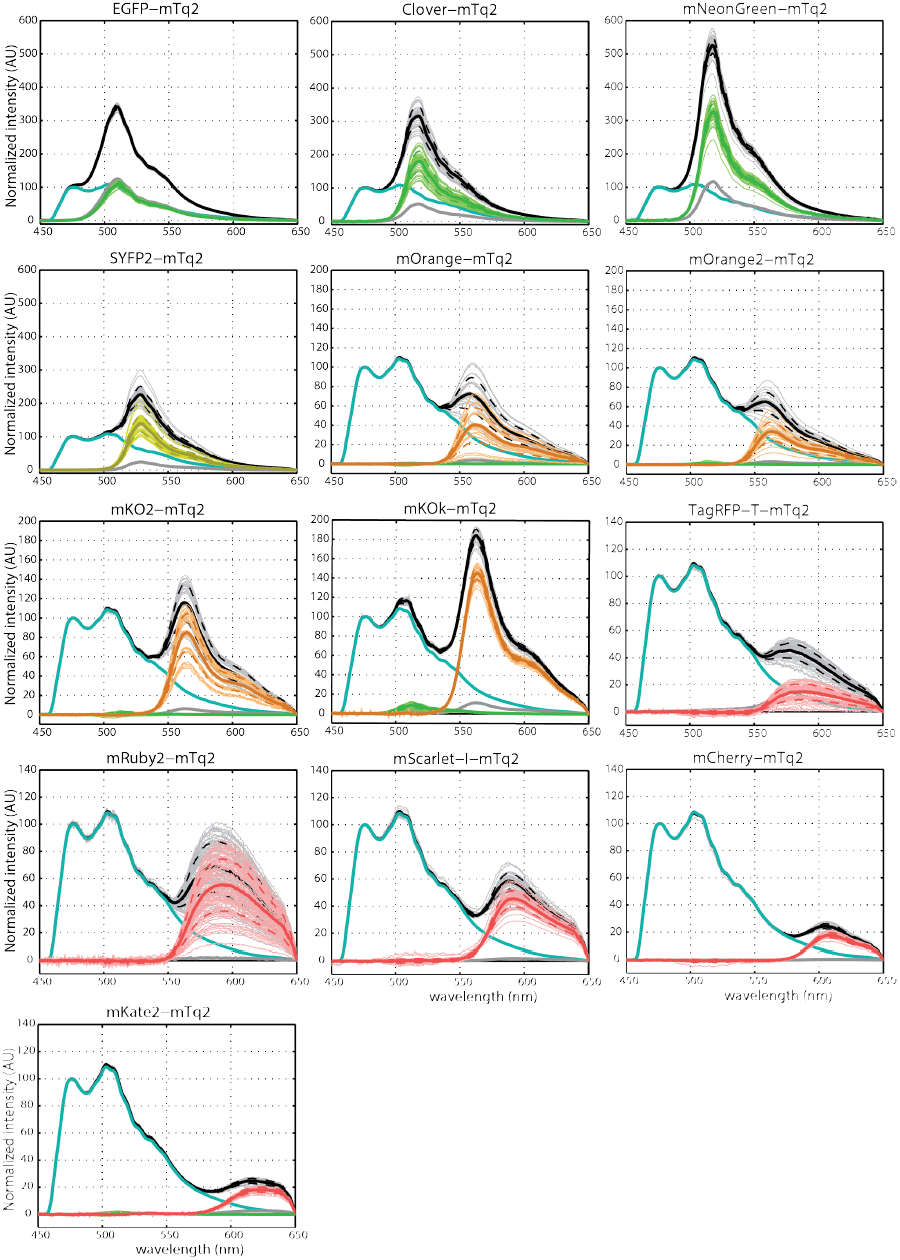
Spectral images of the FRET donor mTurquoise2 fused to different FRET acceptors. The emission spectra of FRET pairs were recorded from single living cells. The sensitized emission component was calculated by unmixing the donor spectrum and the direct acceptor excitation. Black lines represent the FRET-pair spectra. Cyan lines represent the donor emission spectra. Grey lines represent direct acceptor excitation spectra. If orange or red fluorescent proteins show an evident green component, this is represented by a green line. Lines in color of the acceptor emission represent the unmixed sensitized emission. Thick lines show the average emission spectrum, broken lines represent the standard deviations and thin lines show individual measurements. Based on these data the FRET efficiency was calculated (table 3). The number of cells imaged is EGFP *n*=37, Clover *n*=36, mNeonGreen *n*=46, SYFP2 *n*=39, mOrange *n*=24, mOrange2 *n*=22, mKO2 *n*=35, mKO*κ n*=24, TagRFP-T *n*=50, mRuby2 *n*=66, mScarlet-I *n*=47, mCherry *n*=28, mKate2 *n*=27.

As can be inferred from figure 4, there is a large variation in the amount of sensitized emission between the fusion proteins. Overall, the strongest sensitized emission signal is observed for the fusion with mNeonGreen. In the orange spectral class, the fusion with mKO*κ* shows the highest level of sensitized emission and in the red spectral class, we observed relatively high sensitized emission for the fusions with mRuby2 and mScarlet-I. We also note that the cell-to-cell variation differs between FRET pairs. For instance, there is enormous variation between cells in the amount of sensitized emission for the FRET pair with mRuby2. In contrast the amount of sensitized emission for the FRET pair with mScarlet-I is well-defined.

**Table 3.**
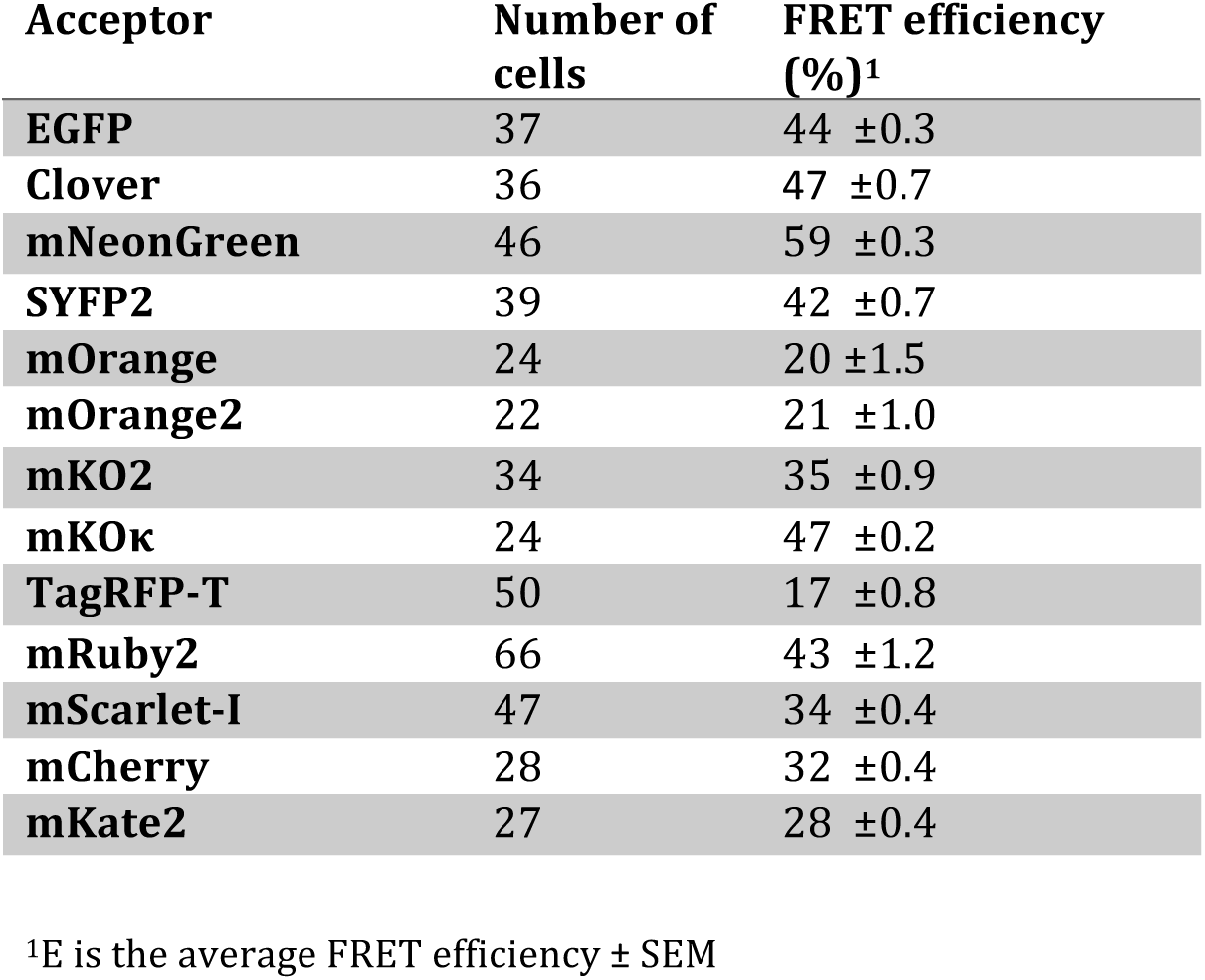
FRET efficiencies of mTurquoise2 paired with the different acceptors, calculated from spectral imaging results (Figure 4).

The differences among the FRET pairs with orange fluorescent proteins were striking. The FRET pair with mKO*κ* showed much stronger sensitized emission than mKO2 which is surprising given the single amino acid difference. In addition, the modest sensitized emission for FRET pairs with mOrange and mOrange2 is unexpected given their high intrinsic brightness (Shaner et al., 2004; Shaner et al., 2008). To further examine the properties of the orange fluorescent proteins, we determined their brightness in cells. We used a previously established assay that measures the fluorescence of transfected cells relative to a quantitatively co-expressed control, in this case mTurquoise2 ^42^. The results show that the brightness in cells is in the order mKO*κ* > mKO2 = mOrange > mOrange2 (supplemental figure S1), showing that mKO*κ* is by far the brightest orange fluorescent protein in cells.

The data depicted in figure 4 was used to calculate the FRET efficiency based on the assumption that every photon emitted by the acceptor stems from a quenched donor photon (see materials and methods). The FRET efficiency value for each FRET pair is listed in table 3. The mTurquoise2-SYFP2 pair showed a FRET efficiency of 42%, in line with the FLIM results. The pair with mNeonGreen as acceptor showed the highest FRET efficiency of 59%, with little cell-to-cell variation. The pair with mKO*κ* shows a relatively high FRET efficiency of 47%. The FRET pair with mRuby2 shows a high FRET efficiency of 43% but this is accompanied by substantial cell-to-cell variability. The FRET efficiency of FRET pairs with mScarlet-I and mCherry is comparable with values of 34% and 32% respectively. Based on the spectral imaging data and R0 values, we prepared an animation of the spectral changes that occur as function of distance for the mTurquoise2-SYFP2 and –mNeonGreen pair (supplemental movie S1) and the mTurquoise2-mCherry and –mScarlet-I pair (supplemental movie S2).

In summary, the FRET efficiencies calculated from the SPIM data are corresponding to the FRET efficiencies calculated from the FLIM data. mNeonGreen shows the highest FRET efficiency and a dominant sensitized emission peak. mKO*κ* shows the highest FRET efficiency and sensitized emission peak of the orange variants. The TagRFP-T and mRuby2 show high cell-to-cell variability. In contrast, mScarlet-I and mCherry show little variation and mScarlet-s I shows a higher amount of sensitized emission than mCherry which is explained by the higher quantum yield of mScarlet-I. Based on the FLIM and SPIM data shown in the previous two paragraphs a selection was made of the most promising FRET acceptors. This selection includes mNeonGreen, mKO*κ*, mRuby2 and mScarlet-I, with as reference commonly used acceptors SYFP2 and mCherry. Out of curiosity for FRET between mTurquoise2 and a far-red acceptor and because FLIM results showed a lifetime decrease comparable to SYFP2, mKate2 was included as well.

### Photostability of FRET pairs

Photostability is a crucial parameter for the reliable and robust detection of FRET, especially in time-lapse imaging. We evaluated the photostability of a selection of fusions with mTurquoise2 that was made based on FRET efficiency. In order to stay close to the purpose of the fluorescent proteins as acceptor in FRET experiments we used the fusion constructs and bleached them under the conditions that are normally used for recording ratiometric FRET data. To determine photostability we continuously excited the donor, while alternatingly measuring donor emission and acceptor emission. The only difference compared to recording FRET data of biosensors is that the photostability measurements are done under continuous illumination instead of 200ms exposure per frame and for a longer duration than usual FRET measurements. The photostability curves for the fusion constructs are depicted in figure 5 and supplemental figure S2. Under these conditions unfused, unquenched mTurquoise2 shows a decrease in intensity over time ^42^. For the tandem fusions, an increase in the CFP channel is observed, which is accompanied by a decrease of acceptor fluorescence. These observations indicate acceptor bleaching, either by FRET or direct excitation, resulting in dequenching of the donor due to diminished FRET.

**Figure 5.**
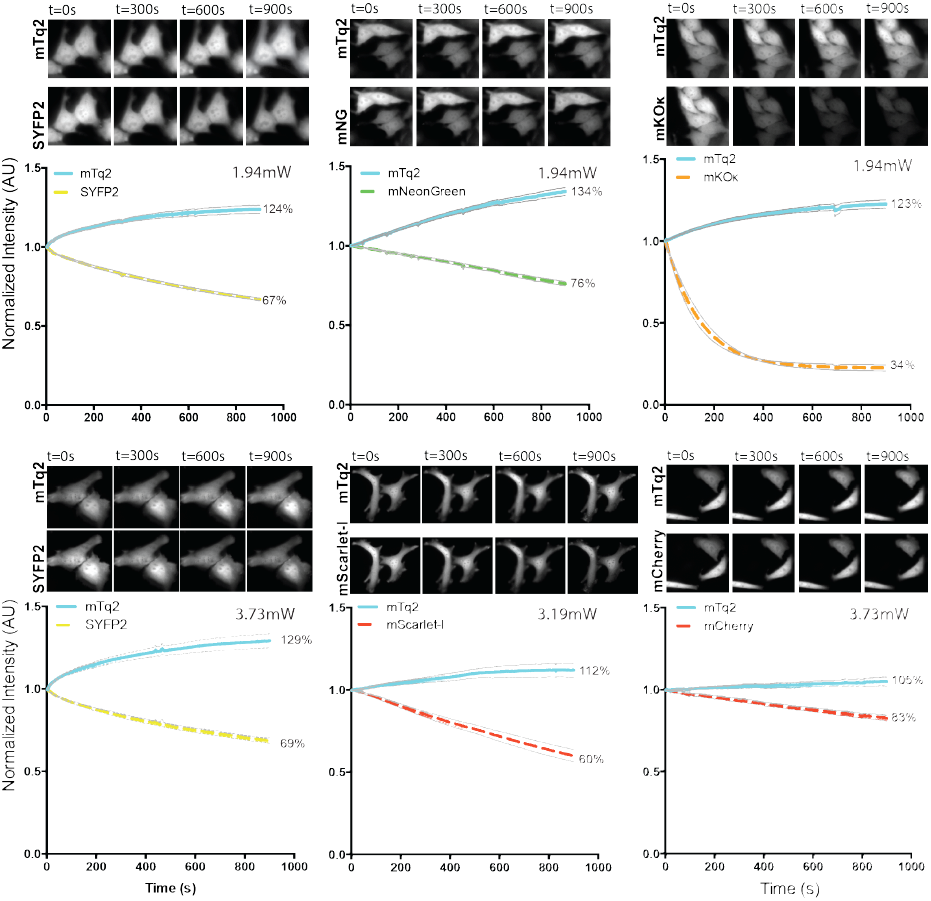
Photostability of tandem pairs during ratiometric FRET measurements. Fusion constructs of mTurquoise2 and acceptor fluorescent protein were used in this experiment. The power is shown in the graphs. The thin lines display the 95% confidence intervals. The photostability of the fusion constructs is shown under continuous illumination with 420nm light for 900s. Images of cells after 0s, 300s, 600s and 900s illumination show the fluorescence intensity. The width of the images are 58.14μm for SYFP2-mTurquoise2 (1.94mW), 87.21μm for mNeonGreen-mTurquoise2, 80.07μm for mKO*κ*-mTurquoise2, 116.28μm for SYFP2-mTurquoise2 (3.73mW), 147.56μm for mScarlet-I-mTurquoise2 and 116.28μm for mCherry-mTurquoise2. For the graph the initial fluorescence intensity was set on 100% and it is stated what percentage of the initial fluorescence is left after 900s illumination. The number of cells imaged is: SYFP2-mTq2 (1.94mW) *n*=23; mNeonGreen-mTq2 *n*=21; mKO*κ*-mTq2 *n*=15; SYFP2-mTq2 (3.73mW) *n*=23; mScarlet-I-mTq2 *n*=11; mCherry-mTq2 *n*=15.

mNeonGreen and SYFP2 are bleached in a similar fashion after 900s continuous illumination, but mNeonGreen bleaches in a more linear fashion while SYFP2 bleaches more rapidly at the start of the experiment. When performing a typical FRET experiment, corresponding to a total exposure time of 48s (240 images of 200ms exposure time),^59^mNeonGreen would be more photostable than SYFP2 (supplemental figure S3). In line with a previous report (Klarenbeek et al., 2015), we observed good photostability of the mTurquoise2-sREACh pair (supplemental figure S2). A striking result is the fast bleaching of mKO*κ* under our conditions. After 48 seconds, which equals a typical FRET measurement, 79% of the initial intensity is left (supplemental figure S3). These results are in line with a previous observation that the related mKO can be photoconverted to a green species by blue light ^15^. When directly exciting mKO*κ* with 570nm light, instead of 420nm, the fluorescent protein is rather photostable (83% of initial intensity left after 900s continuous illumination), and outperforms mKO2 (supplemental figure S4).

mRuby2 shows relatively slow and linear bleaching and after 900s 55% of the initial intensity is left, which is comparable to the photostability of mTurquoise2-SYFP2 with the same excitation power (supplemental figure S2). mKate2 is less photostable than mCherry after 900s of continuous illumination (supplemental figure S2). From figure 5, it can be inferred that the photostability of mScarlet-I is lower than that of mCherry under FRET imaging conditions. Still, under typical conditions for a dynamic FRET experiment (supplemental figure S3), the mScarlet-I hardly loses its intensity.

Recently, a more photostable YFP was reported, generated by one mutation resulting in Y145L ^60^. We mutated SYFP2 and confirmed highly improved photostability (supplementary figure S5).However, next to the reported reduction in brightness, the mTurquoise2-SYFP2(Y145L) pair shows hardly any FRET as compared to the mTurquoise2-SYFP2 pair (supplementary figure S5). In summary, mNeonGreen is a relatively photostable acceptor under FRET ratio imaging conditions. mKO*κ* bleaches rapidly when illuminated with 420nm light and is therefore unfit as FRET acceptor for time-lapse imaging.

### Emission ratio-imaging with novel biosensors to measure heterotrimeric G-protein activation

We used a well-characterized FRET biosensor that measures heterotrimeric G-protein activation to examine how the selection of FRET pairs would perform in terms of dynamic range ^49,59,61^. Since it is of importance to use only monomeric fluorescent proteins for our multimeric membrane located biosensor we evaluated oligomerization by the OSER assay ^38^. A recent thorough analysis of fluorescent proteins that was published during the course of our experiments ^40^, fits largely with our observations. The notable exception is mRuby2, which in our hands shows predominant localization at the Golgi as shown in supplemental figure S6 and documented in more detail recently ^36^. Since this does not represent correct localization, we decided to exclude mRuby2 as acceptor.

The original FRET sensor consists of three subunits that are co-expressed from a single plasmid, including a Gαq tagged with mTurquoise, an untagged Gβ subunit and a Gγ tagged with acceptor ^61^. We modified this plasmid in several ways. First, we replaced mTurquoise by mTurquoise2. The second modification is the removal of the untagged Gβ which turned out to be non-essential (supplementary figure S7). Finally, the Gγ subunit was tagged with the acceptors SYFP2, mNeonGreen, mScarlet-I and mCherry. The resulting plasmid encoded Acceptor-Gγ-IRES-Gαq-mTurquoise2. The localization of the sensor with different acceptor fluorescent proteins is shown in supplementary figure S8.

To examine the FRET response upon activation, we co-expressed the histamine-1 receptor (H1R), which activates the heterotrimeric G-protein, resulting in a loss of FRET. The response is deactivated with the H1R antagonist pyrilamine. As can be inferred from figure 6, the activation of the H1R results in a loss of FRET as inferred from a donor increase and a concomitant acceptor intensity decrease. The sensor with mNeonGreen excels compared to the other acceptors with an increase in donor intensity of up to 30%, when the Gα subunit is activated, while the sensor with SYFP2 showed an increase of approximately 16%. From the acceptor/donor ratio traces it can be concluded that the dynamic range of the sensor with mNeonGreen is higher than that of sensor with SYFP2.

**Figure 6.**
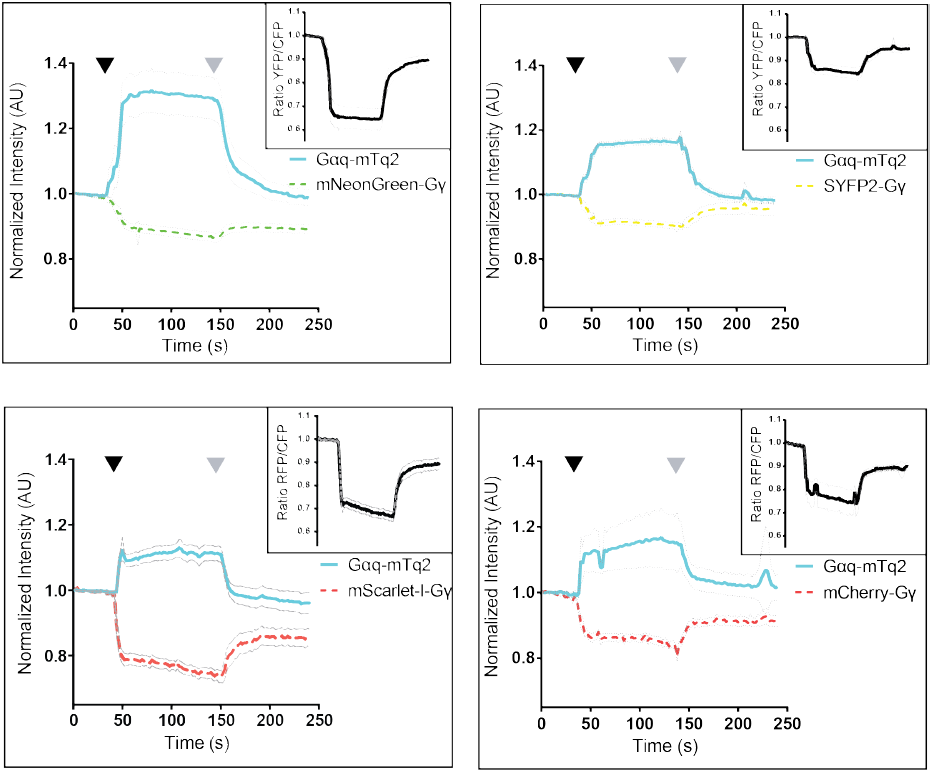
Ratiometric FRET imaging of Gq-activation biosensors equipped with novel FRET pairs. FRET ratioimaging was performed on Hela cells over-expressing the histamine-1 receptor and a FRET biosensor for Gq activation. The blue, solid lines show the mTurquoise2 fluorescence intensity over time, the dashed lines show the acceptor emission level over time. The initial fluorescence intensity is normalized to the average intensity of the first 5 frames. The black graph in a separate upper right window shows the FRET ratio over time. The thin lines indicate the 95% confidence intervals. 100μM histamine was added after 42-50s (black arrowhead) and 10μM pyrilamine was added after 140-150s (grey arrowhead). The number of cells analysed is: Gqsensor-mTq2-mNeonGreen *n*=32 (out of 34 in total), Gqsensor-mTq2-SYFP2 *n*=42 (out of 44 in total), Gqsensor-mTq2-mScarlet-I *n*=24 (out of 26 in total) and Gqsensor-mTq2-mCherry *n*=19 (out of 26 in total)

The same sensor with mCherry as acceptor showed a robust change in FRET. Strikingly, the change in both CFP and RFP fluorescence resulted in a similar dynamic range compared with that of the CFP-YFP variant. The Gq sensor that employs mScarlet-I as the acceptor, shows a more robust decrease of the RFP signal, resulting in a higher dynamic range than the mCherry variant. Often, ratiometric measurements are corrected for CFP bleed-through in the acceptor channel. This is straightforward for the CFP-YFP pair, since the YFP emission is absent in the CFP channel. However, whether this is true for other combinations is unknown. Since chromophore formation of RFP requires several reactions possibly resulting in a fraction of blue or green emitting structures, we expressed RFPs and measured their emission spectra (Supplementary figure S9). We noted different extents of blue/green emission when the RFPs where excited at 436nm, showing that bleedthrough-correction for CFP-RFP pairs is not straightforward. Our data shows that full filter FRET analysis, as reported previously ^36^, is required to calculate the amount of sensitized emission. In summary, employing mNeonGreen as acceptor in the ratiometric FRET sensor for Gq activation yields an improvement of the dynamic range compared to YFP.

### Fluorescence lifetime analysis of heterotrimeric G-protein activation

Since our FLIM analysis showed that sREACh is an efficient FRET acceptor, we evaluated the performance of the G-protein biosensor for FLIM. To this end we constructed a sensor with sREACh as the acceptor. Next, we repeated the GPCR activation/deactivation experiment for sensors incorporating mNeonGreen, SYFP2 or sREACh based sensors with FLIM (Figure 7). All three sensors showed a robust increase in lifetime upon GPCR stimulation, which agrees with the deqeunching of the donor observed by ratio-imaging. The increase in lifetime for the sensor with sREACh of about 0.2ns is in line with previous data ^62^. Addition of pyrilamine, which switches the receptor off, results in a decrease in donor lifetime. Together, these results show that both sREACh and mNeonGreen are suitable acceptors for FLIM in combination with mTurquoise2.

**Figure 7.**
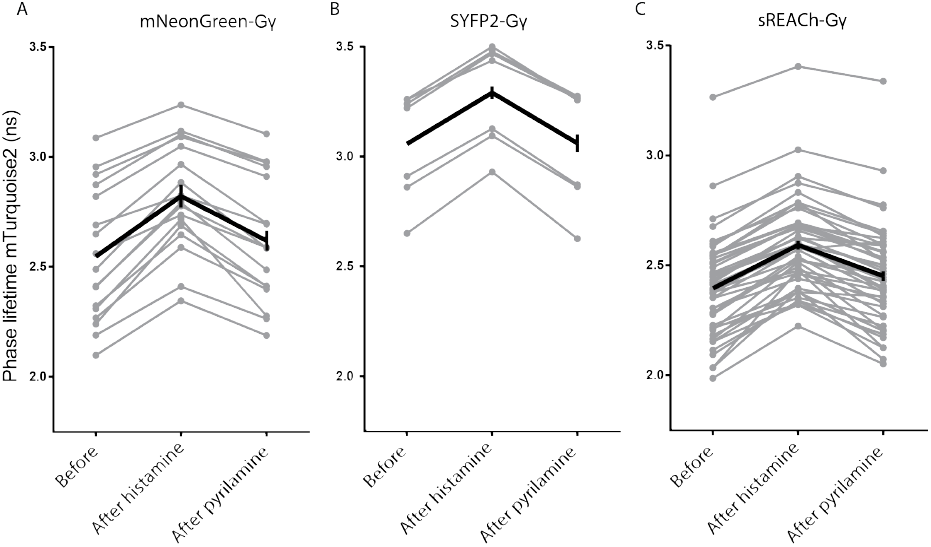
FlIM-FRET of Gq activation biosensors equipped with novel FRET pairs. The fluorescence lifetime of mTurquoise2 was recorded from the biosensor for Gαq activation containing mNeonGreen, SYFP2 or sREACh as FRET acceptor. The phase lifetime was recorded before addition of (ant)agonist, 20-60s after addition of 100μM histamine and 20-60s after addition of 10μM pyrilamine. The changes in phase lifetime are shown in the graphs. The grey lines represent individual cells and the black graph represents the average of which the error bars indicate the 95% confidence intervals. The number of cells used for the graph is for mNeonGreen as FRET acceptor *n*=17 (out of a total of 26 cells), for SYFP2 as FRET acceptor *n*=7 (out of a total of 23 cells) and for sREACh as FRET acceptor *n*=46 (out of a total of 60 cells).

## Discussion

In this study, we have evaluated the performance of FRET pairs consisting of mTurquoise2 as donor and acceptors varying from green to far-red in mammalian cells. In all our experiments the acceptor mNeonGreen consistently showed the highest FRET efficiency and dynamic range, accompanied by strong sensitized emission. This can be explained by the large spectral overlap, high extinction coefficient and high quantum yield of mNeonGreen. It also implies that the maturation efficiency of mNeonGreen is high. Another beneficial feature of mNeonGreen is that it shows increased photostability under FRET conditions relative to SYFP2. However, since photostability is generally power-dependent ^40^, the photostability will differ when experimental conditions are different. We conclude that mTurquoise2-mNeonGreen is the optimal FRET pair for live cell imaging application in mammalian cells and we demonstrate that the mTurquoise2-mNeonGreen pair can be used to generate biosensors with high dynamic range and photostability. The superior performance of mNeonGreen in FRET pairs with mTurquoise2 relative to Clover is surprising, given their equal spectroscopic properties ^40^. We compared the cellular brightness of Clover and mNeonGreen and did not find striking differences (supplemental figure S10). Next, to verify dimerization tendency of mNeonGreen, we performed an OSER assay. The results indicate that mNeonGreen shows no strong tendency to dimerize, in line with previous findings ^40,63^. Therefore, the better performance of mNeonGreen may be explained by better maturation in the context of fusion proteins.

It is of note that the FRET efficiency may be cell-type and certainly will be organism dependent, since the protein maturation can vary in different systems. The set of fluorescent protein fusions, and accompanying controls that we have generated, provides a way to systematically determine the performance of FRET pairs in other biological contexts.

The detection of Gq activation with biosensors based on sREACh or mNeonGreen using FLIM resulted in a similar contrast. Since the blue-shifted emission of mNeonGreen requires a narrow band-pass (BP) filter for exclusive detection of CFP fluorescence in FLIM, the donor emission intensity will be reduced (supplemental figure S11). Therefore, it may be more desirable to use the non-emitting acceptor sREACh for FRET based biosensors that are dedicated for FLIM. Although the properties of the mTurquoise2-mNeonGreen pair are favorable and we demonstrate good performance in an intermolecular FRET sensor, it remains to be determined whether this pair will beat the CFP-YFP or TFP-YFP pair in intramolecular biosensors. Our preliminary data on replacing YFP with mNeonGreen, without any optimization, show that the dynamic range of these sensors is not superior (supplemental figure S12). These preliminary studies suggest that in addition to probe properties, other factors such as linker length, weak homodimerization and probe orientation, determine FRET contrast. Optimization of unimolecular sensors with the mTurquoise2-mNeonGreen pair will benefit from strategies that generate and screen a large number of different variants ^51,64^. Perhaps circular permutation of mTurquoise2 and mNeonGreen would be a viable way to generate high-contrast intramolecular FRET biosensors.

Based on theory, i.e. *R*_*0*_ values, all pairs should display considerable FRET efficiency, but this is not observed in cells for all pairs. We suspect that this is due to inefficient maturation of the acceptor fluorescent protein, affecting the FRET efficiency ^17,58^. Additionally, slight differences in orientation between different acceptors due to small changes in acceptor fluorescent protein length and sequence might also influence the FRET efficiency ^9,65^.

Our photostability analysis of FRET pairs shows a donor increase due to photobleaching of the acceptor. This demonstrates the superior photostability of mTurquoise2 as compared to the employed acceptors. We did not pursue the origin of acceptor photobleaching, which is either due to direct excitation or due to FRET. Regardless of the mechanism, the kinetics of the donor intensity increase will depend on the FRET efficiency of the fluorescent protein pair. In dynamic FRET ratio-imaging, the FRET efficiency can change in time and space and effects of photobleaching will be difficult to correct for. Hence, it is essential for reliable FRET ratio-imaging experiments to avoid photobleaching and to employ the most photostable donor-acceptor pair.

In general, the relatively low quantum yields of the red fluorescent proteins result in a low level of sensitized emission, which is a disadvantage for the acquisition of ratiometric FRET data. Although mRuby2 combines efficient FRET with a high sensitized emission, several properties argue against the use in FRET experiments combined with mTurquoise2. First, the fusion with CytERM shows Golgi localization, rather than the ER localization, making it impossible to reliably determine monomeric behavior (supplemental figure S6). Second, a large fraction of blue fluorescence of mRuby2 is detected in the CFP emission band (supplemental figure S9). Finally, the FRET efficiency is variable between cells, which can likely be attributed to variable maturation ^17^ or photochromism ^36^. Similar cell-to-cell variation is observed for tagRFP-T. The recently developed monomeric RFP, mScarlet-I, with good maturation and high quantum yield has optimal properties for ratiometric FRET. Our results show that mScarlet-I exhibits high sensitized emission amongst the RFPs tested when paired with mTurquoise2. We conclude that mScarlet-I is the preferred acceptor in the red part of the spectrum since (i) it shows consistent FRET in cells, with substantial sensitized emission, (ii) it has little blue fluorescence when excited at 436nm, (iii) it shows good photostability when excited at the CFP excitation wavelength and (iv) it is monomeric ^36^. In addition, we demonstrate that it can be used in a biosensor to report on the activation of a GPCR. In summary, the results obtained in this study point out that mTurquoise2-mNeonGreen is an optimal FRET pair for ratiometric detection of cellular processes with genetically encoded intermolecular FRET based biosensors.

## Materials and Methods

### Cloning / plasmid construction

All fluorescent proteins (FPs) that were used as FRET acceptor, were cloned in clontech-style C1 mammalian expression vectors and RSET bacterial expression vectors with flanking AgeI and BsrgI restriction sites. RSET bacterial expression vectors were used for protein production and isolation while the C1 vectors were used to construct the fusion constructs (figure 2).

TagRFP-T was made by introducing the S158T point mutation into tagRFP ^43^. Clover and mRuby2 were derived from a plasmid obtained from addgene (#40255), mCherry-C1 and mOrange-C1 were previously described ^15^. mOrange2 was a kind gift of M. Ouyang ^66^. mKO2 was a kind gift of R.N. Day ^67^. mKO*κ* ^68^ was obtained by introducing the point mutation M176F (Fw: 5’-GGCAATCACAAATGCCAATTCAAGACTACTTACAAGGCG-3’ Rv: 5’-CGCCTTGTAAGTAGTCTTGAATTGGCATTTGTGATTGCC-3’) in the mKO2 coding sequence. sREACh was obtained from addgene (plasmid #21949) ^69^ mKate2 was a kind gift of D.M. Chudakov ^70^ mNeonGreen was a kind gift of N.C. Shaner ^63^. mScarlet-I has been reported by Bindels et al ^36^.All tandem fusions were based on the SYFP2-mTurquoise2 construct that was previously described ^42^; addgene #60493). The SYFP2 in the plasmids SYFP2-mTurquoise2 was replaced by the acceptor fluorescent protein of interest cut from the clontech-style C1 plasmids using NdeI/Kpn2I restriction enzymes.

The brightness of fluorescent proteins was analyzed using tandem FP constructs with a T2A linker resulting in equal expression of two fluorescent proteins ^71^. These constructs were produced by cutting the SYFP2-mTurquoise2 with BamhI/Kpn2I and inserting two hybridized oligonucleotides (5min, 95°C) ^72^ (Fw: 5-ccggagagggcagaggaagtcttctaacatgcggtgacgtggaggagaatcccggccctgt-3’; Rv: 5’-gatccagggccgggattctcctccacgtcaccgcatgttagaagacttcctctgccctct-3’), resulting in SYFP2-T2A-mTurquoise2. mTurquoise2 is used as reference in the brightness assay to correct for protein concentration. The SYFP2 is replaced by a FP of which the brightness is to be characterized, cut from the clontech-style C1 plasmids using NdeI/Kpn2I restriction enzymes.

The multimeric biosensors for Gq activation are based on the published FRET biosensor that is encoded on a single plasmid; Gβ1-2A-YFP-Gγ2-IRES-Gαq-mTurquoiseΔ6 ^61^. In order to make the Gq activation biosensors with the desired FRET pairs, first, mTurquoiseΔ6 was exchanged for mTurquoise2Δ6 in a pcDNA3.1 vector containing the Gαq-mTurquoiseΔ6 sequence, where the fluorescent protein is inserted at the 125^th^ amino acid residue of the Gαq sequence. A PCR was performed on a clontech-style C1 vector containing the mTurquoise2 sequence using primers ’-TTGAGGATCCAAGCGGAGGCGGAGGCAGCATGGTGAGCAAGGGCG-3’and Rv 5’-GTATATGCCGAGAGTGATCCCGGC-3’. The PCR product and the pcDNA3.1 Gαq-mTurquoiseΔ6 vector were both digested with BamHI and SnaBI (PCR product digested with only BamHI since half the SnaBI site is present in the reverse primer, which can be directly ligated in the SnaBI cut vector) and the digested mTurquoise2Δ6 PCR product was ligated in the Gαq pcDNA3.1 vector. Subsequently, the Gαq-mTurquoise2Δ6 vector and the Gβ1-2A-YFP-Gγ2-IRES-Gαq-mTurquoiseΔ6 sensor were both digested with BamHI and EcoRI. Then, the Gαq-mTurquoise2Δ6 was ligated in the sensor replacing the original Gαq-mTurquoiseΔ6, leading to Gβ1-2A-YFP-Gγ2-IRES-Gαq-mTurquoise2Δ6 sensor. In order to exchange the YFP-Gγ2 in the sensor for other acceptor fluorescent proteins (FP), first, acceptor FP-Gγ2 fusions were constructed. A PCR was performed on clontech-style C1 vector containing the Gγ2 sequence using primers Fw 5’-AGCTGTACATGGCCAGCAACAACACC-3’ and Rv 5’-TCTACAAATGTGGTATGGC-3’. The Gγ2 PCR product and clontech-style C1-FP plasmids were digested with BsrGI and SacII and the Gγ2 sequence is ligated behind the fluorescent protein in the clontech-style C1 vector. Next, these acceptor FP-Gγ2 fusions and the Gβ1-2A-YFP-Gγ2-IRES-Gaq-mTurquoise2Δ6 sensor were digested with NheI and SacII and the FP-Gγ2 sequence was ligated into Gβ1-2A-YFP-Gγ2-IRES-Gaq-mTurquoise2Δ6 replacing the original Gβ1-2A-YFP-Gγ2. Finally, this resulted in a pcDNA vector encoding acceptorFP-Gγ2-IRES-Gαq-mTurquoise2Δ6.

The intramolecular FRET sensors for RhoA activation are based on the published sensor (DORA-RhoA) ^73,74^. First a PCR is performed on a clontech-style C1-Tq2(206A) (nTq2) plasmid using the primers Fw 5’-AACGGATCCGTGAGCAAGGGCGAGG-3’ and Rv 5’-AGCGCTAGCCCCGGCGGCGGTCAC-3’. The PCR product and the original RhoA sensor were digested with BamHI and NheI and the nTq2 was ligated in the sensor construct replacing the original donor Cerulean3. A BglII restriction site is introduced in the sensor plasmid, behind the acceptor FP sequence and simultaneously the original acceptor is swapped for mNeonGreen via overlap-extension PCR ^75,76^. The CRs were performed on the clontech-style C1 plasmid containing mNeonGreen using primerAfirst P Fw 5’-CTACCGGTGCCACCATG-3’ and primerB Rv 5’-CTCGATGTTAGATCTGAGTCCGGACTTGTACA-3’ and on the RhoA activation sensor containing the correct donor FP using primerC Fw 5’-CTCAGATCTAACATCGAGGAAGCACAAAAG-3’ and primerD Rv 5’-TGCACGTGTATACAGCTGTGC-3’.The second PCR was performed on a mix of both PCR products using primerA and primerD. This second PCR product and the RhoA sensor are digested with AgeI and HindIII and the PCR product containing the BglII restriction site and mNeonGreen is ligated into the sensor. To swap the acceptor from mNeonGreen to SYFP2 a PCR was performed on a clontech-style C1 vector containing SYFP2 using Fw 5’-CTACCGGTGCCACCATG-3’ and Rv 5’-TCTACAAATGTGGTATGGC-3’ and both PCR product and sensor (containing mNeonGreen) were digested with AgeI and BglII and the SYFP2 is ligated in the sensor replacing mNeonGreen.

The intramolecular FRET sensors for calcium are based on the published Twitch2B sensor addgene (#49531) ^64^. In order to swap the fluorescent proteins the calcium binding domain and the acceptor FP were transferred to a RSET bacterial expression plasmid, using SphI and EcoRI, resulting in RSET-Minimal Calcium Binding Domain-cpmCitrine. A PCR was performed on a RSET vector containing mTurquoise2 using the primers: Fw 5’-TAATACGACTCACTATAGGG-3’ and Rv 5’-GGTCATGCATGCGGGCGGCGGTCACGAAC-3’. The PCR product and RSET-Minimal Calcium Binding Domain -cpmCitrine vector were both digested with NcoI and SphI and mTurquoise2 was inserted prior to the calcium binding domain sequence. A mutagenesis PCR is performed on the Twitch2B RSET plasmid introducing a XhoI restriction site (by introducing 3 nucleotides) using primers Fw 5’-CCCATCTACCCCGAGCTCGAGATGGGTGGGGTC-3’ and Rv 5’-GACCCCACCCATCTCGAGCTCGGGGTAGATGGG-3’. Then a PCR is performed on a clontech-style C1 vector containing either mNeonGreen or SYFP2 using Fw 5’-GAGATCTCGAGATGGTGAGCAAGGGCG-’3 and Rv 5’-GAGCTGAATTCTCACTTGTACAGCTCGTCCATGC-’3. The PCR product and mutagenized Twitch2B RSET plasmid are both digested with XhoI and EcoRI and ligated, exchanging the acceptor FP for mNeonGreen or SYFP2. With NheI and EcoRI the whole sensor module is transferred from the RSET vector to a clontech-style C1 vector for mammalian expression. Plasmids generated in this study will be available through addgene at http://www.addgene.org/Dorus_Gadella/

### Spectroscopy of purified fluorescent proteins

His6-tagged proteins were produced in E.coli and purified on Hisbind resin (Novagen, Darmstadt, Germany), according to Bindels et.al. ^77^. After elution by imidazole the proteins were dialyzed 2x against 20mM Tris. Spectral measurements were done in 20mM Tris ^15^, unless indicated otherwise. Absorption spectra were recorded on a Libra S70 double-beam spectrophotometer (Biochrom) ^42^ Emission spectra were recorded on a Perkin Elmer LS55 fluorimeter. Emission spectra were recorded with following settings: mKO*κ* ex525nm, slit 5nm; em530-750nm slit 5nm; scan speed 150nm/min; pmt 750V. mOrange ex530nm, slit 5nm; em540-750nm slit 2.5nm; scan speed 150nm/min; pmt 750V. mOrange2 ex530nm, slit 5nm; em540-750nm slit 2.5nm; scan speed 150nm/min; pmt 760V. mKO2 ex520nm, slit 5nm; em535-750nm slit 5nm; scan speed 150nm/min; pmt 750V. mNeonGreen ex460nm, slit 5nm; em470-675nm slit 5nm; scan speed 150nm/min; pmt 760V. Clover ex465nm, slit 5nm; em475-650nm slit 5nm; scan speed 150nm/min; pmt 760V. sREACh ex505nm, slit 5nm; em515-675nm slit 5nm; scan speed 150nm/min; pmt 810V. mCherry, mScarlet-I, mRuby2, tagRFP-T and mKate2 ex540nm slit 2.5nm; em550-800nm slit 2.5nm; scan speed 150nm/min in PBS (50mM PO_4_, 136mM NaCl, 2.7mM KCl, pH7.4).

Emission spectra were corrected for instrument response factors after calibration with emission spectra of established fluorophores. The emission spectra of SYFP2 and EGFP were acquired previously ^18,78^.

The *R*_*0*_ values were calculated as described previously ^11,15^.

### Cell culture and transfection

HeLa cells (CCL-2, American Tissue Culture Collection; Manassas,VA, USA) were cultured in Dulbecco’s modified Eagle’s medium (DMEM) (Gibco, cat# 61965–059) supplemented with 10% fetal bovine serum (Invitrogen, cat# 10270-106), 100U/ml penicillin and 100μg/ml streptomycin at 37°C in 7% CO2. For microscopy experiments cells were grown on 24mm Ø round coverslips, 0.13 - 0.16mm thick (Menzel, cat# 360208) to 50% confluency and transfected with 500ng plasmid DNA, 1μL Lipofectamin 2000 (Invitrogen, cat# 11668–019), 2μl Polyethylenimine (PEI) (1mg/ml) in EtOH, or 4.5μl PEI (1mg/ml) in water (pH 7.3) and 100μl OptiMEM (Gibco, cat# 31985-047) per 35mm Ø dish holding a 24mm Ø coverslip. Two days after transfection the coverslip was mounted in a cell chamber (Attofluor, Invitrogen). Microscopy medium (20mM HEPES (pH = 7.4), 137mM NaCL, 5.4mM KCl, 1.8mM CaCl2, 0.8mM MgCl2 and 20mM glucose) was added to the coverslip in the cell chamber. The OSER assay, Ratiometric FRET, bleaching and brightness experiments are performed at 37°C.

### Fluorescence lifetime imaging microscopy

Fluorescence lifetime imaging was performed using the wide-field frequency domain approach on a home-build instrument ^79^ using a RF-modulated AOM and a RF-modulated image intensifier (Lambert Instruments II18MD) coupled to a CCD camera (Photometrics HQ) as detector. A 40x objective (Plan NeoFluar NA 1.3 oil) was used for all measurements. The modulation frequency was set to 75.1MHz. At least twelve phase images with an exposure time of 20–100ms seconds were acquired in a random recording order to minimize artifacts due to photobleaching ^80^. A picoquant directly modulated diode laser was used for excitation at 442nm, passed onto the sample by 455dclp dichroic and emission light was filtered by a BP480/40 emission filter. When imaging with GFP as FRET acceptor, a second emission filter BP447/60 was combined with the BP480/40 filter (supplemental figure S11). Each by a reference FLIM measurement is calibrated measurement of the reflected laser light using a modified filter cube ^79^ for correcting the phase and modulation drift of the excitation light. The reference is calibrated by averaging five FLIM measurements of cells expressing mTurquoise2 (mTq2), which has a known phase lifetime of 3.8ns and a modulation lifetime of 4.0ns ^42^. This extra calibration corrects for path-length differences and possible opticsrelated reflections that are different between the FLIM and reference measurements. At least twelve phase sequences were acquired from each sample. From the phase sequence, an intensity (DC) image, phase and modulation lifetime images were calculated ^81^using Matlab macros. Alternatively, we performed the fluorescence lifetime measurements with a Nikon Eclipse Ti-E inverted microscope equipped with a LIFA system (Multi-Led illumination and LI2CAM; Lambert Instruments). The modulated 446nm LED excitation light passed through a 448/20 excitation filter (FF01-448/20, Semrock), reflected towards the sample by a 442nm dichroic mirror (Di02-R442, Semrock) and focused using a 60x objective (Nikon, CFI Plan Apochromat NA 1.4 oil, MDR01605). The emission was filtered by a BP482/20 (FF01-482/25, Semrock). The LI-FLIM software (Li-FLIM 1.223 Lambert Instruments) recorded 18 phase steps (with three times averaging) in pseudorandom order at a frequency of 40MHz. Erythrosin B (198269, Sigma-Aldrich) dissolved in ddH2O was used as reference dye (fluorescence lifetime 0.086 ns; ten times averaging for reference stack). After background subtraction and 3×3 blurring, the lifetimes were calculated by the LI-FLIM software.

The FRET efficiency *E* was calculated according to: *E*= (1- (*τ*_*DA*_ /*τ*_*D*_)) ^*^ 100%, in which *τ*_*DA*_ is the fluorescence lifetime of the donor in presence of the acceptor and *τ*_*D*_ is the fluorescence lifetime of the donor in absence of the acceptor. Since frequency domain FLIM yields a phase lifetime and a modulation lifetime, the FRET efficiency can be calculated based on both ^15^.

For the fluorescence lifetime analysis of heterotrimeric G-protein activation the same methods were used to measure the fluorescence lifetime before adding 100uM histamine, 20-60s after adding histamine and 20-60s after adding 10uM pyrilamine.

### Spectral imaging microscopy of FRET pairs

Spectral imaging of living cells was performed with hardware as described ^82^, two days after transfection using an imaging spectrograph-CCD detector.

For each cell transfected with a construct of interest a spectral image was acquired using donor excitation at 436/20nm, an 80/20 (transmission/reflection) dichroic and a 460LP (long-pass) emission filter. Subsequently a spectral image was acquired using acceptor excitation without exciting the donor. For EGFP, mNeonGreen, Clover and SYFP2 (green/yellow) excitation at 500/20nm and for detection a BP534/20 filter was used, for mKO2, mKO*κ*, mOrange, mOrange2 (orange) excitation at 500/20nm and for detection a 530LP filter was used, and for mScarlet-I, mRuby2, mCherry, TagRFP-T and mKate2 (red) excitation at 546/10nm and for detection a 590LP filter was used. Using a custom made Matlab script, cells were selected from the spectral images and each sample spectrum obtained with donor excitation settings was normalized to the peak intensity of the spectrum obtained using acceptor excitation settings. In general, the donor excitation setting also leads to direct excitation of the acceptor. Using cells transfected with an acceptor only construct the direct acceptor excitation contribution could be estimated. The donor-only spectrum was obtained by using cells transfected with mTurquoise2. Prior to unmixing, all spectra were aligned and the wavelength axis was calibrated. From each sample spectrum, *F*(*λ*), the direct acceptor excitation spectrum, *F*_*A*_ (*λ*), was subtracted in order to remove the contribution of direct acceptor excitation. For the green/yellow acceptors, the donor component, *F*_*D*_(*λ*) and the sensitized emission component, *F*_*S*_ (*λ*) were obtained from the spectrum with linear regression using the donor-only spectrum and acceptor-only spectrum, both obtained with donor excitation settings. In this case the whole wavelength range (450-650nm) was used. For the orange and red variants with a green component the donor and green contribution, *F*_*G*_(*λ*) were obtained by unmixing the sample spectrum in the wavelength range of 450-525 using the donor-only spectrum and the EGFP-only spectrum. The sensitized emission was then obtained by subtracting the unmixed donor and green component from the spectrum. For red variants without a discernable green component in the spectra a similar approach was used, but now only using the donor-only spectrum applied to a wavelength range of 450-500nm. All sample and unmixed spectra *F(λ), F*_*D*_ (*λ*), *F*_*S*_ (*λ*) and *F*_*G*_ (*λ*) were subsequently normalized to the first peak value of the donor. Using the unmixed donor and sensitized emission spectrum the apparent energy transfer, *E*_*D*_ can be estimated using the following equation ^83^:

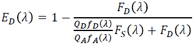

Where *Q*_*D*_ and *Q*_*A*_ denote the quantum yields of the donor and acceptor respectively, and *f*_*D*_ (λ) and *f*_*D*_ (λ) denote corrected and area normalized reference spectra of the donor and acceptor respectively. The numerator represents the quenched donor and the denominator represents the total donor emission reconstructed using the ratio of quantum yields and reference spectra. In principle the *E*_*D*_ value should be the same for each wavelength, hence it should follow a flat line, however at the edge of spectra this can deviate and therefore a flat region was selected and a weighted average was calculated using weights *w* (λ) = *f*_*D*_ (λ) *f*_*A*_ (λ), hence *E*_*D*_ = Ʃ_*i*_ *w* (λ_*i*_)*E*_*D*_ (λ_*i*_)/ Ʃ_*i*_ *w* (λ_*i*_)

### Photostability

Photostability of fluorescent proteins in fusion constructs was measured on a wide-field fluorescence microscope (Axiovert 200 M; Carl Zeiss GmbH) equipped with a xenon arc lamp with monochromator (Cairn Research, Faversham, Kent, UK). Measurements were performed under continuous illumination for 900s with 420nm light (slit width 30nm) to excite mTurquoise2. Supplemental figure S3 shows the photobleaching results for the first 48s continuous illumination (corresponding to the total illumination time during a FRET experiment with 200ms exposure time and 121 time frames) of the same experiments as shown in figure 5 and supplemental figure S2. The power was measured at the 20x objective (Zeiss LD-A-plan 20x Air/0,30 ph1 ∞) using a coherent power meter (FM Fieldmaster Power Energy Meter, 0210-761-99). Each 4s, fluorescence intensity of FRET donor and acceptor was recorded with an exposure time of 200ms using a 40x objective (oil-immersion Plan-Neo-fluor 40×/1.30; Carl Zeiss GmbH). mTurquoise2 emission was detected with a BP470/30 filter, GFP/YFP emission was detected with a BP535/30 filter and OFP/RFP emission was detected with a BP620/60 filter ^49^. Image analysis was done in ImageJ. After subtraction of background signal, the mean fluorescence intensity of the cells was calculated for each time point.

### Ratiometric FRET measurements

Zeiss FRET ratio-imaging was performed on a wide-field fluorescence microscope (Axiovert 200 M; Carl GmbH) ^49^ equipped with a xenon arc lamp with monochromator (Cairn Research, Faversham, Kent, UK) for 240s and with a time interval of 2s. The fluorescence intensity of the donor and acceptor were recorded with an exposure time of 200ms per image using a 40x objective (oilimmersion Plan-Neo-fluor 40×/1.30; Carl Zeiss GmbH). HeLa cells were used expressing Gq-sensors, comprising Gαq-mTq2 and acceptor FP-Gγ, and histamine-1 receptor-2A-mCherry ^59^ or oin the case of orange or red acceptors in the Gq-sensor untagged histamine-1 receptor ^49^. Fluorophores were excited with 420nm light (slit width 30nm), mTq2 emission was detected with the BP470/30 filter, GFP/YFP emission was detected with the BP535/30 filter and OFP/RFP emission was detected with BP620/60 filter by turning the filter wheel (supplemental figure S11). After 42-50s HeLa cells were stimulated with 100µM (final concentration) histamine (Sigma-Aldrich) and after 140-150s 10µM (final concentration) pyrilamine (mepyramine) (Sigma-Aldrich) was added as antagonist. The curves were corrected for shifts in time point of adding drugs. The curves were normalized to the average intensity of the first 5 frames that were recorded. ImageJ was used to perform a background correction and calculation of mean intensity of each cell for each time point. Cells that did not show a visible response were not used for the analysis. The total number of cells imaged and the number of cells analyzed (“the responders”) are indicated in the figure legends.

For the FRET measurements using the RhoA activation biosensors or the calcium sensors, the same time lapse, filter settings, exposure times and analysis methods are used. For the calcium sensor, HeLa cells are stimulated with 100μM Histamine at t=44s (black arrow) and at t= 150s with 10μg/ml Ionomycin (Cayman chemical #10004974) (gray arrow). For the RhoA sensor, cells are stimulated with 100μM Histamine at t=44s (black arrow) and antagonized at t= 150s with 10μM Pyrilamine (gray arrow). In these cells, a GEFT-mCherry construct was overexpressed next to the FRET sensor ^84^.

### Brightness analysis

Cells were transfected with Tandem FP constructs containing a T2A linker, mTurquoise2 as reference and a fluorescent protein of interest. Cells expressing two separate FPs in equal amounts were imaged on a widefield fluorescence microscope (Axiovert 200 M; Carl Zeiss GmbH) equipped with a xenon arc lamp with monochromator (Cairn Research, Faversham, Kent, UK), using a 40x objective (oil-immersion Plan-Neo-fluor 40×/1.30; Carl Zeiss GmbH). Orange FPs were excited with 510nm light and emission was detected with a BP572/25 filter. As reference, mTurquoise2 was excited with 420nm light and emission was detected with a BP470/30 filter. Clover and mNeonGreen are excited with 500nm light and emission was detected with a BP535/30 filter. To prevent cross excitation, reference mTurquoise2 was excited with 405nm light and emission was detected with a BP470/30 filter. After subtraction of background signal, the mean fluorescence intensity of the cells was calculated. The fluorescence intensity of the protein of interest relative to the fluorescence intensity of the reference mTurquoise2 reveals the relative brightness of the protein of interest ^42^.

## Acknowledgments

We thank the members of our lab for their continuous interest and support of this project.

## Author contributions

M.M. D.S.B. and J.G. performed experiments and analyzed data. M.M. and J.G. and wrote the manuscript. N.C.S. contributed essential reagents and data M.P. developed image analysis methods and assisted with the data analysis. T.W.J.G. assisted with experimental design and interpretation of data. All authors approved the final manuscript.

## References

1. Chudakov, D. M., Matz, M. V, Lukyanov, S. & Lukyanov, K. A. Fluorescent proteins and their applications in imaging living cells and tissues. Physiol Rev 90, 1103–1163 (2010).

2. Tsien, R. Y. the Green Fluorescent. Proteins 67, 509–44 (1998).

3. Miyawaki, A. & Niino, Y. Molecular Spies for Bioimaging — Fluorescent Protein-Based Probes. Mol. Cell 58, 632–643 (2015).

4. Chudakov, D. M., Lukyanov, S. & Lukyanov, K. A. Fluorescent proteins as a toolkit for in vivo imaging. Trends Biotechnol 23, 605–613 (2005).

5. Gadella Jr, T. W. J., van der Krogt, G. N. & Bisseling, T. GFP-based FRET microscopy in living plant cells. Trends Plant Sci. 4, 287–291 (1999).

6. Miyawaki, A. Development of probes for cellular functions using fluorescent proteins and fluorescence resonance energy transfer. Annu. Rev. Biochem. 80, 357–73 (2011).

7. Pollok, B. A. & Heim, R. Using GFP in FRET-based applications. Trends Cell Biol. 9, 57–60 (1999).

8. Piston, D. W. & Kremers, G. J. Fluorescent protein FRET: the good, the bad and the ugly. Trends Biochem. Sci. 32, 407–14 (2007).

9. Jares-Erijman, E. A. & Jovin, T. M. FRET imaging. Nat Biotechnol 21, 1387–1395 (2003).

10. Pietraszewska-Bogiel, A. & Gadella Jr, T. W. J. FRET microscopy: from principle to routine technology in cell biology. J. Microsc. 241, 111–8 (2011).

11. Wu, P. G. & Brand, L. Resonance energy transfer: methods and applications. Anal. Biochem.218, 1–13 (1994).

12. Hamers, D., van Voorst Vader, L., Borst, J. W. & Goedhart, J. Development of FRET biosensors for mammalian and plant systems. Protoplasma 251, 333–47 (2014).

13. Mehta, S. & Zhang, J. Reporting from the field: genetically encoded fluorescent reporters uncover signaling dynamics in living biological systems. Annu. Rev. Biochem. 80, 375–401 (2011).

14. Okumoto, S., Jones, A. & Frommer, W. B. Quantitative imaging with fluorescent biosensors.Annu. Rev. Plant Biol. 63, 663–706 (2012).

15. Goedhart, J., Vermeer, J. E., Adjobo-Hermans, M. J., van Weeren, L. & Gadella Jr, T. W. J. Sensitive Detection of p65 Homodimers Using Red-Shifted and Fluorescent Protein-Based FRET Couples. PLoS One 2, e1011 (2007).

16. Van der Krogt, G. N. M., Ogink, J., Ponsioen, B. & Jalink, K. A comparison of donor-acceptor pairs for genetically encoded FRET sensors: application to the Epac cAMP sensor as an example. PLoS One 3, e1916 (2008).

17. Scott, B. L. & Hoppe, A. D. Optimizing fluorescent protein trios for 3-Way FRET imaging of protein interactions in living cells. Sci. Rep. 5, 10270 (2015).

18. Kremers, G. J., Goedhart, J., van Munster, E. B. & Gadella Jr, T. W. J. Cyan and yellow super fluorescent proteins with improved brightness, protein folding, and FRET Förster radius. Biochemistry 45, 6570–80 (2006).

19. Cubitt, A. B., Woollenweber, L. a & Heim, R. Understanding Structure-Function Relationships in the Aequorea victoria Green Fluorescent Protein. Methods Cell Biol. 58, 19–30 (1998).

20. Siemering, K. R., Golbik, R., Sever, R. & Haseloff, J. Mutations that suppress the thermosensitivity of green fluorescent protein. Curr. Biol. 6, 1653–1663 (1996).

21. Fukuda, H., Arai, M. & Kuwajima, K. Folding of green fluorescent protein and the Cycle3 mutant. Biochemistry 39, 12025–12032 (2000).

22. Patterson, G. H., Knobel, S. M., Sharif, W. D., Kain, S. R. & Piston, D. W. Use of the green fluorescent protein and its mutants in quantitative fluorescence microscopy. Biophys. J. 73, 2782–90 (1997).

23. Cormack, B. P., Valdivia, R. H. & Falkow, S. FACS-optimized mutants of the green fluorescent protein (GFP). Gene 173, 33–38 (1996).

24. Nagai, T. et al. A variant of yellow fluorescent protein with fast and efficient maturation for cell-biological applications. Nat. Biotechnol. 20, 87–90 (2002).

25. Wachter, R. M., Watkins, J. L. & Kim, H. Mechanistic diversity of red fluorescence acquisition by GFP-like proteins. Biochemistry 49, 7417–27 (2010).

26. Miyawaki, A., Shcherbakova, D. M. & Verkhusha, V. V. Red fluorescent proteins: Chromophore formation and cellular applications. Curr. Opin. Struct. Biol. 22, 679–688 (2012).

27. Griesbeck, O., Baird, G. S., Campbell, R. E., Zacharias, D. a & Tsien, R. Y. Reducing the environmental sensitivity of yellow fluorescent protein. Mechanism and applications. J. Biol. Chem. 276, 29188–94 (2001).

28. Rekas, A., Alattia, J.-R., Nagai, T., Miyawaki, A. & Ikura, M. Crystal structure of venus, a yellow fluorescent protein with improved maturation and reduced environmental sensitivity. J. Biol. Chem. 277, 50573–8 (2002).

29. Verkhusha, V. V. & Lukyanov, K. A. The molecular properties and applications of Anthozoa fluorescent proteins and chromoproteins. Nat. Biotechnol. 22, 289–96 (2004).

30. Baird, G. S., Zacharias, D. A. & Tsien, R. Y. Biochemistry, mutagenesis, and oligomerization of DsRed, a red fluorescent protein from coral. Proc. Natl. Acad. Sci. U. S. A. 97, 11984–9 (2000).

31. Zacharias, D. A. Partitioning of Lipid-Modified Monomeric GFPs into Membrane Microdomains of Live Cells. Science (80-). 296, 913–916 (2002).

32. Vinkenborg, J. L., Evers, T. H., Reulen, S. W. a, Meijer, E. W. & Merkx, M. Enhanced sensitivity of FRET-based protease sensors by redesign of the GFP dimerization interface. Chembiochem 8, 1119–21 (2007).

33. Lindenburg, L. H. et al. Quantifying stickiness: thermodynamic characterization of intramolecular domain interactions to guide the design of förster resonance energy transfer sensors. Biochemistry 53, 6370–81 (2014).

34. Nguyen, A. W. & Daugherty, P. S. Evolutionary optimization of fluorescent proteins for intracellular FRET. Nat. Biotechnol. 23, 355–60 (2005).

35. Campbell, R. E. et al. A monomeric red fluorescent protein. Proc. Natl. Acad. Sci. U. S. A. 99, 7877–82 (2002).

36. Bindels, D. S. et al. mScarlet: a bright monomeric red fluorescent protein for cellular imaging. Nat. Methods 14, 53–56 (2016).

37. Pédelacq, J.-D., Cabantous, S., Tran, T., Terwilliger, T. C. & Waldo, G. S. Engineering and characterization of a superfolder green fluorescent protein. Nat. Biotechnol. 24, 79–88 (2006).

38. Costantini, L. M., Fossati, M., Francolini, M. & Snapp, E. L. Assessing the Tendency of Fluorescent Proteins to Oligomerize Under Physiologic Conditions. Traffic 13, 643–649 (2012).

39. Costantini, L. M. et al. A palette of fluorescent proteins optimized for diverse cellular environments. Nat. Commun. 6, 7670 (2015).

40. Cranfill, P. J. et al. Quantitative assessment of fluorescent proteins. Nat. Methods 13, 557– 562 (2016).

41. Van Munster, E. B., Kremers, G. J., Adjobo-Hermans, M. J. W. & Gadella Jr, T. W. J. Fluorescence resonance energy transfer (FRET) measurement by gradual acceptor photobleaching. J. Microsc. 218, 253–262 (2005).

42. Goedhart, J. et al. Structure-guided evolution of cyan fluorescent proteins towards a quantum yield of 93%. Nat. Commun. 3, 751 (2012).

43. Shaner, N. C. et al. Improving the photostability of bright monomeric orange and red fluorescent proteins. Nat. Methods 5, 545–551 (2008).

44. Cody, C. W., Prasher, D. C., Westler, W. M., Prendergast, F. G. & Ward, W. W. Chemical structure of the hexapeptide chromophore of the Aequorea green-fluorescent protein. Biochemistry 32, 1212–1218 (1993).

45. Swaminathan, R., Hoang, C. P. & Verkman, A. S. Photobleaching recovery and anisotropy decay of green fluorescent protein GFP-S65T in solution and cells: cytoplasmic viscosity probed by green fluorescent protein translational and rotational diffusion. Biophys. J. 72, 1900–7 (1997).

46. Cubitt, A. B. et al. Understanding, improving and using green fluorescent proteins. Trends Biochem. Sci. 20, 448–455 (1995).

47. Greenbaum, L., Rothmann, C. & Lavie, R. Green Fluorescent Protein Photobleaching : a Model for Protein Damage by Endogenous and Exogenous Singlet Oxygen. Biol. Chem. 381, 1251– 1258 (2000).

48. Nagai, T., Yamada, S., Tominaga, T., Ichikawa, M. & Miyawaki, A. Expanded dynamic range of fluorescent indicators for Ca(2+) by circularly permuted yellow fluorescent proteins. Proc. Natl. Acad. Sci. U. S. A. 101, 10554–9 (2004).

49. Adjobo-Hermans, M. J. et al. Real-time visualization of heterotrimeric G protein Gq activation in living cells. BMC Biol. 9, 32 (2011).

50. Klarenbeek, J., Goedhart, J., van Batenburg, A., Groenewald, D. & Jalink, K. Fourth-generation epac-based FRET sensors for cAMP feature exceptional brightness, photostability and dynamic range: characterization of dedicated sensors for FLIM, for ratiometry and with high affinity. PLoS One 10, e0122513 (2015).

51. Fritz, R. D. et al. A versatile toolkit to produce sensitive FRET biosensors to visualize signaling in time and space. Sci. Signal. 6, rs12 (2013).

52. Komatsu, N. et al. Development of an optimized backbone of FRET biosensors for kinases and GTPases. Mol. Biol. Cell 22, 4647–56 (2011).

53. Shimozono, S. & Miyawaki, A. Engineering FRET constructs using CFP and YFP. Methods Cell Biol. 85, 381–93 (2008).

54. Peroza, E. A., Boumezbeur, A.-H. & Zamboni, N. Rapid, randomized development of genetically encoded FRET sensors for small molecules. Analyst 140, 4540–8 (2015).

55. Schifferer, M. & Griesbeck, O. A dynamic FRET reporter of gene expression improved by functional screening. J. Am. Chem. Soc. 134, 15185–15188 (2012).

56. Ohta, Y. et al. Nontrivial Effect of the Color-Exchange of a Donor/Acceptor Pair in the Engineering of Förster Resonance Energy Transfer (FRET)-Based Indicators. ACS Chem. Biol. 11, 1816–22 (2016).

57. Goedhart, J., Hink, M. A. & Jalink, K. An introduction to fluorescence imaging techniques geared towards biosensor applications. Methods in Molecular Biology 1071, 17–28 (2014).

58. Shaner, N. C. et al. Improved monomeric red, orange and yellow fluorescent proteins derived from Discosoma sp. red fluorescent protein. Nat. Biotechnol. 22, 1567–72 (2004).

59. van Unen, J. et al. Quantitative Single-Cell Analysis of Signaling Pathways Activated Immediately Downstream of Histamine Receptor Subtypes. Mol. Pharmacol. 90, 162–176 (2016).

60. Bogdanov, A. M. et al. Turning On and Off Photoinduced Electron Transfer in Fluorescent Proteins by π-Stacking, Halide Binding, and Tyr145 Mutations. J. Am. Chem. Soc. 138, 4807– 17 (2016).

61. Goedhart, J. et al. Quantitative Co-Expression of Proteins at the Single Cell Level – Application to a Multimeric FRET Sensor. PLoS One 6, e27321 (2011).

62. Raspe, M. et al. siFLIM: single-image frequency-domain FLIM provides fast and photon-efficient lifetime data. Nat. Methods 13, 501–4 (2016).

63. Shaner, N. C. et al. A bright monomeric green fluorescent protein derived from Branchiostoma lanceolatum. Nat. Methods 10, 407–409 (2013).

64. Thestrup, T. et al. Optimized ratiometric calcium sensors for functional in vivo imaging of neurons and T lymphocytes. Nat. Methods 11, 175–82 (2014).

65. Shimozono, S. et al. Concatenation of cyan and yellow fluorescent proteins for efficient resonance energy transfer. Biochemistry 45, 6267–6271 (2006).

66. Ouyang, M. et al. Simultaneous visualization of protumorigenic Src and MT1-MMP activities with fluorescence resonance energy transfer. Cancer Res. 70, 2204–2212 (2010).

67. Sun, Y. et al. Characterization of an orange acceptor fluorescent protein for sensitized spectral fluorescence resonance energy transfer microscopy using a white-light laser. J. Biomed. Opt. 14, 54009 (2009).

68. Tsutsui, H., Karasawa, S., Okamura, Y. & Miyawaki, A. Improving membrane voltage measurements using FRET with new fluorescent proteins. Nat. Methods 5, 683–5 (2008).

69. Murakoshi, H., Lee, S. J. & Yasuda, R. Highly sensitive and quantitative FRET-FLIM imaging in single dendritic spines using improved non-radiative YFP. Brain Cell Biol. 36, 31–42 (2008).

70. Shcherbo, D. et al. Far-red fluorescent tags for protein imaging in living tissues. Biochem. J.418, 567–574 (2009).

71. Kim, J. H. et al. High cleavage efficiency of a 2A peptide derived from porcine teschovirus-1 in human cell lines, zebrafish and mice. PLoS One 6, 1–8 (2011).

72. Goedhart, J. & Gadella Jr, T. W. J. Analysis of oligonucleotide annealing by electrophoresis in agarose gels using sodium borate conductive medium. Anal. Biochem. 343, 186–187 (2005).

73. van Unen, J. et al. Plasma membrane restricted RhoGEF activity is sufficient for RhoA-mediated actin polymerization. Sci. Rep. 5, 14693 (2015).

74. Pertz, O., Hodgson, L., Klemke, R. L. & Hahn, K. M. Spatiotemporal dynamics of RhoA activity in migrating cells. Nature 440, 1069–72 (2006).

75. Horton, R. M., Hunt, H. D., Ho, S. N., Pullen, J. K. & Pease, L. R. Engineering hybrid genes without the use of restriction enzymes : gene splicing by overlap extension sequences ; frequency of errors ; exon ; intron ; mosaic fusion protein ; mouse histocompatibility genes). 77, 61–68 (1989).

76. Heckman, K. L. & Pease, L. R. Gene splicing and mutagenesis by PCR-driven overlap extension. Nat. Protoc. 2, 924–32 (2007).

77. Bindels, D. S. et al. in Fluorescence Spectroscopy and Microscopy: Methods and Protocols (eds. Engelborghs, Y. & Visser, J. W. G. A.) 371–417 (Humana Press, 2014). doi:10.1007/978-1-62703-649-8_16

78. Kremers, G. J., Goedhart, J., Van Den Heuvel, D. J., Gerritsen, H. C. & Gadella Jr, T. W. J. Improved green and blue fluorescent proteins for expression in bacteria and mammalian cells. Biochemistry 46, 3775–3783 (2007).

79. van Munster, E. B. & Gadella Jr, T. W. J. phiFLIM: a new method to avoid aliasing in frequency-domain fluorescence lifetime imaging microscopy. J. Microsc. 213, 29–38 (2004).

80. van Munster, E. B. & Gadella Jr., T. W. Suppression of photobleaching-induced artifacts in frequency-domain FLIM by permutation of the recording order. Cytom. A 58, 185–194 (2004).

81. van Munster, E. B. & Gadella Jr, T. W. J. Fluorescence Lifetime Imaging Microscopy (FLIM). Advances in Biochemical Engineering/Biotechnology 95, 143–175 (2005).

82. Vermeer, J. E. M., Van Munster, E. B., Vischer, N. O. & Gadella Jr, T. W. J. Probing plasma membrane microdomains in cowpea protoplasts using lipidated GFP-fusion proteins and multimode FRET microscopy. J. Microsc. 214, 190–200 (2004).

83. Wlodarczyk, J. et al. Analysis of FRET Signals in the Presence of Free Donors and Acceptors. Biophys. J. 94, 986–1000 (2008).

84. van Unen, J. et al. Kinetics of recruitment and allosteric activation of ARHGEF25 isoforms by the heterotrimeric G-protein Gαq. Sci. Rep. 6, 36825 (2016).

85. Lam, A. J. et al. Improving FRET dynamic range with bright green and red fluorescent proteins. Nat. Methods 9, 1005–12 (2012).

86. Ganesan, S., Ameer-Beg, S. M., Ng, T. T. C., Vojnovic, B. & Wouters, F. S. A dark yellow fluorescent protein (YFP)-based Resonance Energy-Accepting Chromoprotein (REACh) for Förster resonance energy transfer with GFP. Proc. Natl. Acad. Sci. U. S. A. 103, 4089–94 (2006).

87. Sakaue-Sawano, A. et al. Visualizing Spatiotemporal Dynamics of Multicellular Cell-Cycle Progression. Cell 132, 487–498 (2008).

